# Picomolar affinity antagonist and sustained signaling agonist peptide ligands for the adrenomedullin and calcitonin gene-related peptide receptors

**DOI:** 10.1101/2020.02.28.970301

**Authors:** Jason M. Booe, Margaret L. Warner, Augen A. Pioszak

**Author notes:** Corresponding author Augen A. Pioszak, Ph.D.

## Abstract

The calcitonin receptor-like G protein-coupled receptor (CLR) mediates adrenomedullin (AM) and calcitonin gene-related peptide (CGRP) actions including vasodilation, cardioprotection, and nociception. Receptor activity-modifying proteins (RAMP1-3) determine CLR ligand selectivity through an unresolved mechanism. CLR-RAMP complexes are drug targets, but short AM and CGRP plasma half-lives limit their therapeutic utility. We used combinatorial peptide library and rational design approaches to probe selectivity determinants and develop short AM and CGRP variants with ∼1000-fold increased receptor extracellular domain affinities. Binding and structural studies explained the increased affinities and defined roles for AM Lys46 and RAMP modulation of CLR conformation in selectivity. In longer scaffolds that also bind the CLR transmembrane domain the variants generated picomolar affinity antagonists, one with an estimated 12.5 hr CGRP receptor residence time, and **<u>s</u>**ustained **<u>s</u>**ignaling agonists ss-AM and ss-CGRP. This work clarifies the RAMP-modulated ligand selectivity mechanism and provides AM and CGRP variants with promise as long-acting therapeutics.

## Introduction

The peptides adrenomedullin (AM) and calcitonin gene-related peptide (CGRP) have overlapping and distinct roles in human physiology and pathophysiology^1, 2^. Their actions are mediated by the calcitonin receptor-like receptor (CLR), which is a class B G protein-coupled receptor (GPCR) that is an important drug target. Both peptides exhibit vasodilator activity and cardioprotective actions^3, 4^. AM has crucial roles in cardiac and lymphatic development^5^, and in adults it promotes endothelial barrier integrity and embryo implantation^6, 7^. AM showed beneficial effects in treating heart attack and inflammatory bowel disease in pilot clinical trials^8, 9^, and holds promise for heart failure, sepsis, and infertility^6, 10, 11^. CGRP is a neuropeptide involved in pain transmission and neurogenic inflammation and it has a crucial role in migraine headache pathogenesis^2, 12, 13^. Monoclonal antibodies and a small molecule that antagonize CGRP signaling recently obtained regulatory approval for migraine^12^. CGRP antagonists also showed promise for treating necrotizing fasciitis in a mouse model^14^, and the CGRP agonist inhibited mucosal HIV transmission in a cell model^15^. Both AM and CGRP antagonists may be of value for treating various cancers^16, 17^.

There is substantial potential for AM and CGRP agonists and antagonists as novel therapeutics, however, the short plasma half-lives (∼20 min.)^18, 19^ of AM and CGRP limit their therapeutic utility. This is particularly problematic for indications where the agonists are the desired drugs because there are no small molecule CLR agonist alternatives. AM or CGRP analogs with PEGylation, acylation, or other modifications to enhance plasma half-life have been developed^20–22^, but these have reduced signaling potencies. As an alternative or in addition to prolonging circulatory half-life, there is growing recognition of the power of increasing receptor residence time to achieve efficacious long-lasting drug action^23, 24^. Although typically applied to inhibitors, this concept can also be of value for GPCR agonists^25, 26^. Long residence time AM and CGRP analogs might overcome their short plasma half-lives, but to our knowledge no such analogs have been reported.

AM and CGRP binding to CLR is controlled by three receptor activity-modifying proteins (RAMP1-3) that heterodimerize with CLR^27^. RAMP1 favors CGRP binding, giving the CGRP receptor, whereas RAMP2 and −3 favor AM binding, giving the AM_1_ and AM_2_ receptors, respectively. A related peptide, adrenomedullin 2/intermedin (AM2/IMD), binds the receptors somewhat non-selectively. CLR and the RAMPs each have extracellular (ECD) and transmembrane (TMD) domains that associate with their counterparts in the heterodimer. Peptide binding to CLR follows the class B GPCR “two-domain” model in which the C-terminal half of the peptide binds the ECD and the N-terminal half binds and activates the TMD^28^. Soluble RAMP-CLR ECD fusion proteins exhibited peptide-binding preferences similar to the intact receptors, but with the lower binding affinities expected from lost peptide-TMD contacts^29, 30^. Crystal structures of C-terminal fragments of a CGRP analog or AM2/IMD bound to RAMP1-CLR ECD and AM bound to RAMP2-CLR ECD showed that the peptides bind a common site on CLR, adopt relatively unstructured conformations defined by a β-turn structural element, and have minimal RAMP contacts^30, 31^. A cryo-EM structure of the intact CGRP receptor with bound CGRP and heterotrimeric G_s_ showed that the N-terminal half of CGRP occupied the CLR TMD with an α-helical conformation and revealed an absence of CGRP-RAMP1 contact outside of the ECD complex^32^.

Understanding how RAMPs determine CLR ligand selectivity is important because this system is a model for accessory membrane protein modulation of GPCR pharmacology and knowledge of the mechanism will aid design of selective therapeutics. The ECD complex is a major selectivity determinant resulting in part from peptide-RAMP contacts^30, 31^. The C-terminal peptide residues AM Y52 and CGRP F37 anchor in a pocket on the CLR ECD that is augmented on one side by the RAMP subunit. AM Y52 forms a hydrogen bond (H-bond) with RAMP2 E101 that is not available in RAMP1 and CGRP F37 makes hydrophobic contacts with RAMP1 W84 that RAMP2 cannot provide. Unfortunately, beyond this the mechanism is unresolved. In the crystal structure a second AM residue, K46, contacts RAMP2, but it is unclear if this contributes to selectivity because it also has an intramolecular packing role that complicates interpretation of mutagenesis data. In addition, subtle CLR ECD conformational differences observed in the structures with RAMP1 and RAMP2 hinted at an allosteric role for the RAMPs^31^, but clear evidence that this is an important component of the selectivity mechanism is lacking.

We previously used rational design to develop truncated ECD-binding AM and CGRP variants with increased affinities by introducing substitutions designed to stabilize the β-turn^33^. Here, we used a combinatorial peptide library approach^34^ to identify novel AM substitutions that further enhance ECD affinity and we present a high-resolution crystal structure that explains the enhanced affinity of a library-identified variant. Results from the library screen and new rationally designed AM and CGRP variants allowed us to define important roles for AM K46 and RAMP modulation of CLR ECD conformation in selectivity. Incorporating key variants in longer peptide scaffolds that also bind the CLR TMD yielded picomolar affinity AM and CGRP antagonists and long-acting, sustained signaling AM and CGRP agonists. These results clarify the selectivity mechanism and provide a suite of novel peptides with immediate value as pharmacological tools and future promise as long residence time therapeutics.

## Results

### Positional scanning-synthetic peptide combinatorial library (PS-SPCL) screen for AM

A PS-SPCL^34^ was designed (see Methods) to identify novel affinity-increasing substitutions in AM and to probe selectivity determinants. The scaffold was the minimal ECD-binding AM(37-52) fragment including the affinity-enhancing Q50W substitution identified by rational design^33^. The 5 positions chosen for substitution were S45, K46, I47, S48 and Y52 (Fig. 1b). S45, K46, and Y52 were chosen because our previous rational design effort indicated that substitutions at these positions could enhance affinity and/or alter selectivity as exemplified by the S45W/K46L/Q50W/Y52F variant that had altered preference and increased affinity for RAMP1-CLR ECD^33^ (Fig. 1a). I47 and S48 were chosen as two new positions not previously explored. The AM PS-SPCL, comprising 95 mixtures and nearly 2.5 million unique theoretical peptides, was screened for binding to the three purified RAMP-CLR ECD complexes (Fig. 1a-c) in a fluorescence polarization (FP) competition binding assay (Fig. 1d).

**Figure 1.**
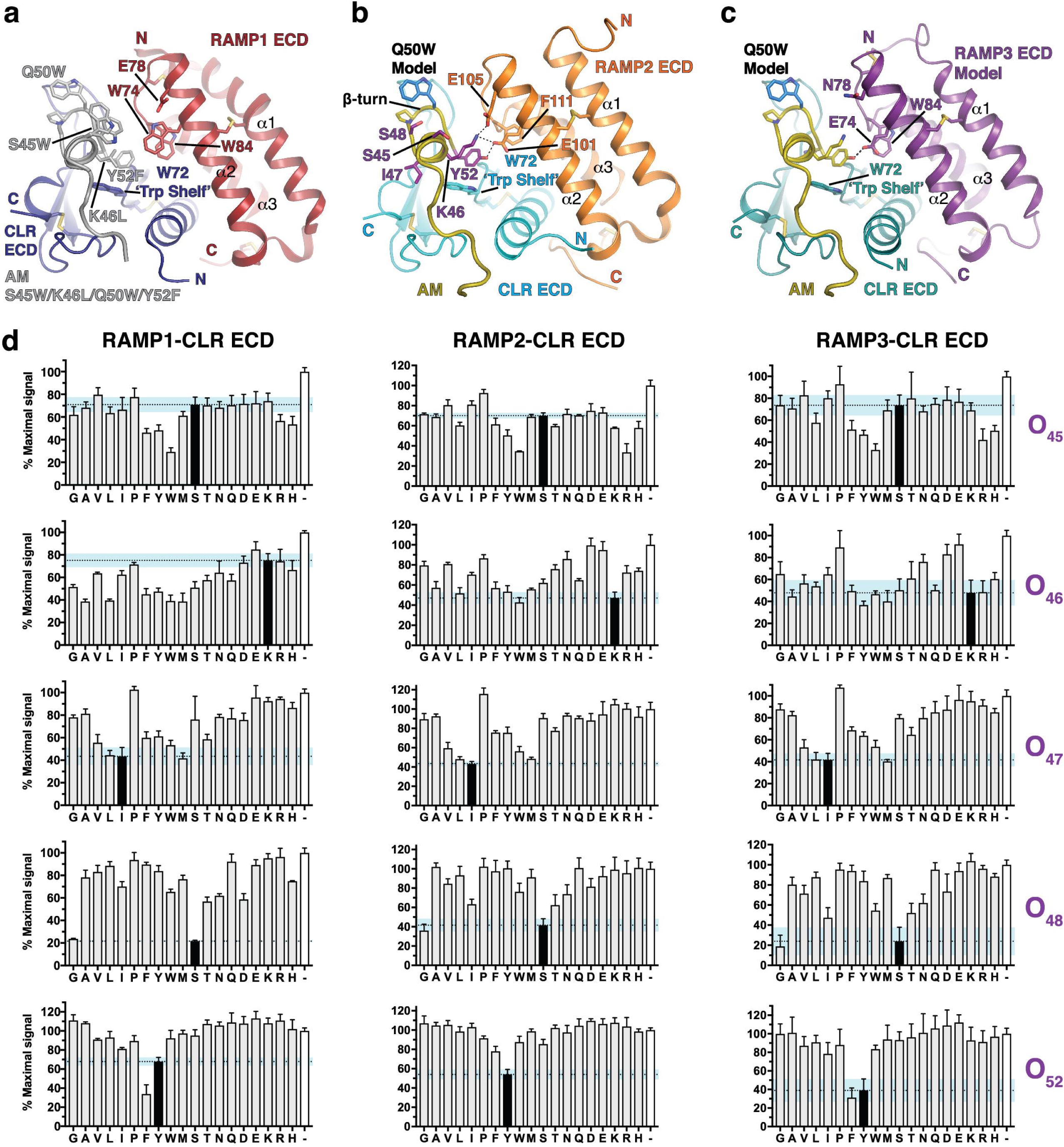
AM(37-52) Q50W PS-SPCL screen at purified MBP-RAMP-CLR ECD fusion proteins. (a) Crystal structure of MBP-RAMP1-CLR ECD with a rationally designed altered selectivity and enhanced affinity AM variant [PDB 5V6Y]. (b) Crystal structure of MBP-RAMP2-CLR ECD with AM [PDB 4RWF]. (c) Homology model of the RAMP3-CLR ECD complex with AM^31^. In (a) and (b) MBP is omitted for clarity. (d) Competition FP assays with the AM PS-SPCL mixtures. Black bars indicate the wild-type residue at the indicated position while the white bars represent no competitor control. Data are mean composite of two independent replicates. Error is shown as standard deviation (SD) with the blue shaded region representing one SD of wild-type residue.

Satisfyingly, the library screen confirmed our prior rational design results. W at position 45 increased affinity at all three complexes and L at position 46 improved binding with RAMP1. F at position 52 diminished binding with RAMP2 while it enhanced binding with RAMP1 and had no effect at the RAMP3 complex (Fig. 1d).

Importantly, the library screen also identified novel and unexpected substitutions. At position 45, F, Y, R, or H also increased affinity, with R perhaps favoring RAMP2/3. Unexpectedly, the library revealed that L at position 46 was tolerated with RAMP2 and indicated that a range of hydrophobic or small polar residues at this position improved binding with RAMP1. Notably, G at position 46 improved binding with RAMP1 and decreased binding with RAMP2/3. These results strongly suggested a significant role for AM K46 in receptor selectivity. For the positions not previously explored, L and M were tolerated at position 47 and G worked surprisingly well at position 48 (Fig. 1d).

### RAMP-CLR ECD complex binding affinities of defined AM variants

The large number of affinity-enhancing substitutions identified made it impractical to test all possible combinations in defined AM variants. Instead, guided by modeling we chose to further examine the effects of the library-identified S45R, K46W/A/G, and S48G substitutions in ten new defined AM(37-52) variants, several of which also included the previous rationally designed K46L, Q50W, or Y52F substitutions. S45R was chosen because it might make ionic interactions with E101 and/or E105 in RAMP2 (Fig. 1b), or E74 in RAMP3 (Fig. 1c). The substitutions at position 46 were chosen to test both small and large residues. S48G was chosen because S48 forms intramolecular H-bonds that appear to stabilize the AM β-turn and α-helical turn^31^, so it was surprising that loss of this side chain did not diminish binding.

First we characterized the affinities of five prior rational design variants^33^ for the purified RAMP-CLR ECD complexes in the competition FP assay (Fig. 2a and Supplementary Table 1). This was done because our previous study lacked the RAMP3-CLR ECD complex and the RAMP1- and −2-CLR ECD complexes that were used were produced in *E. coli* and therefore lacked N-glycosylation. We have since shown that N-glycosylation of the CLR ECD increases peptide-binding affinity^35^. Measuring binding at the three N-glycosylated ECD complexes indicated that Q50W and S45W enhanced affinity from the micromolar into the nanomolar range and K46L and Y52F altered selectivity to favor RAMP1 and disfavor RAMP2 (Fig. 2a).

**Figure 2.**
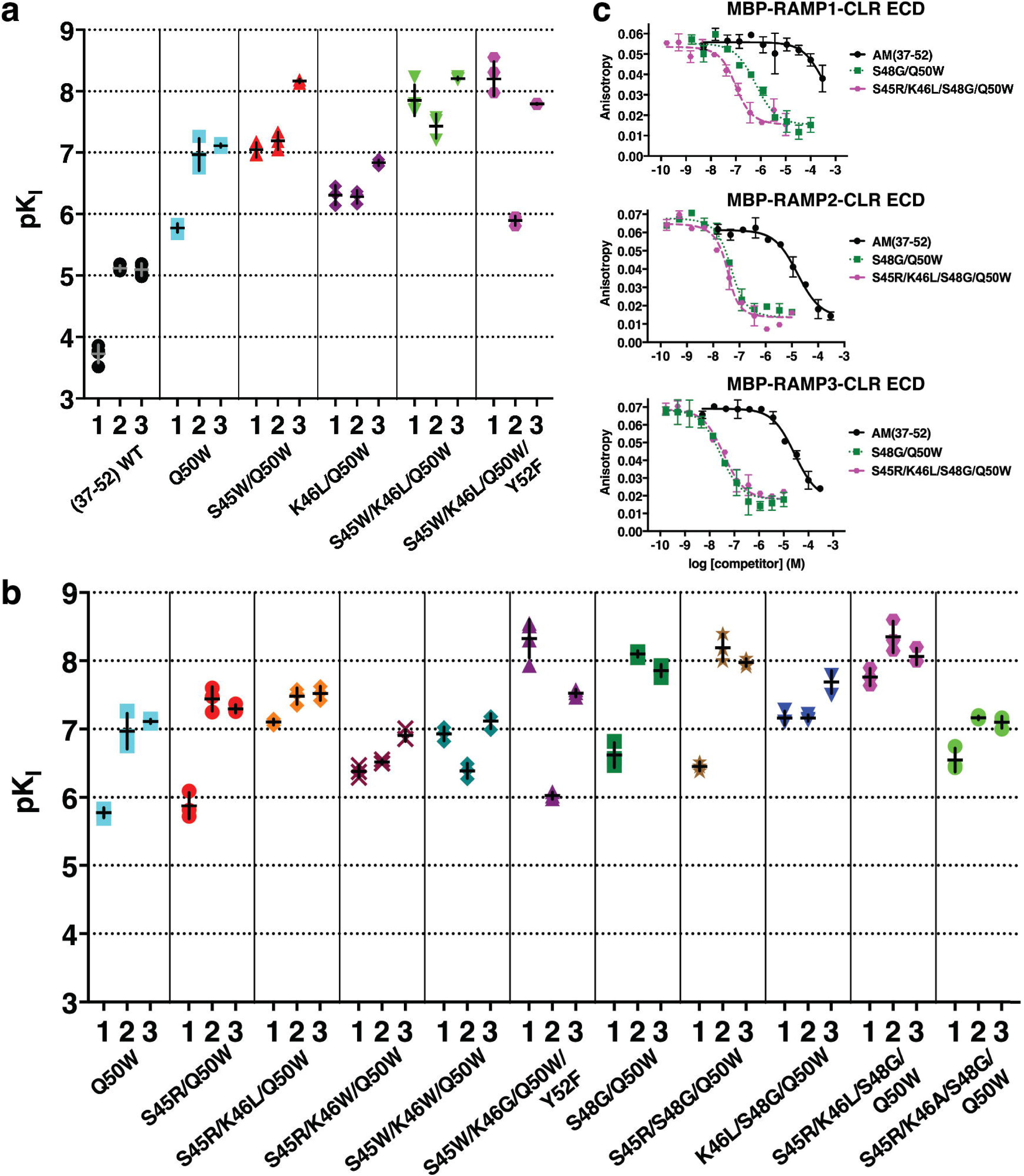
Binding of defined AM(37-52) variants to purified MBP-RAMP-CLR ECD fusion proteins. (a, b) Scatter plot of pK_I_ values from competition FP assays for (a) previously reported, rationally designed AM variants^33^ or (b) new AM variants incorporating library-identified substitutions. Numbers below the x-axis signify the RAMP1, −2, or −3 complexes. (c) Representative competition binding curves for two key variants compared to wild-type. See Supplementary Table 1 for pK_I_ values with SEM, selectivity comparisons, and associated statistical analyses.

Next we examined the binding of new variants that incorporated the library-identified substitutions (Fig. 2b and Supplementary Table 1). S45R had little effect in the S45R/Q50W double mutant as compared to Q50W, however, with K46L in the S45R/K46L/Q50W triple mutant S45R increased affinity ∼5-fold at RAMP1/3 and 17-fold at RAMP2 (as compared to K46L/Q50W). A large Trp at position 46 in the S45R/K46W/Q50W and S45W/K46W/Q50W variants yielded somewhat non-selective peptides with lower affinities, whereas a small Gly at position 46 in the S45W/K46G/Q50W/Y52F quadruple mutant yielded a peptide with altered preference and enhanced affinity for RAMP1-CLR ECD equivalent to our prior rationally designed S45W/K46L/Q50W/Y52F variant. The latter result was quite surprising because we had thought that the bulky Leu at position 46 was needed for selectivity towards RAMP1^33^.

The S48G substitution in the S48G/Q50W double mutant enhanced affinity ∼6-fold at RAMP1/3 and 13-fold at RAMP2 as compared to Q50W (Fig. 2b and Supplementary Table 1). Adding S45R to give the S45R/S48G/Q50W triple mutant had no effect, whereas adding K46L favored RAMP1 and disfavored RAMP2 in the K46L/S48G/Q50W variant. The quadruple mutant S45R/K46L/S48G/Q50W exhibited single digit nanomolar affinities for the RAMP2/3 complexes and double digit nanomolar affinity for the RAMP1 complex. Comparing this to the two prior triple mutants indicated that S45R enhanced affinity in the presence of K46L. Ala at position 46 in the S45R/K46A/S48G/Q50W quadruple mutant decreased affinity ∼10-fold at each of the complexes as compared to the K46L-containing version. Representative FP competition binding curves for the S48G/Q50W and S45R/K46L/S48G/Q50W variants highlight their ∼3 orders of magnitude increased affinities (Fig. 2c).

### Structural basis for enhanced RAMP2-CLR ECD affinity of AM S45R/K46L/S48G/Q50W

To provide insights into how S45R, S48G, and Q50W enhanced affinity, and why S45R only did so in the presence of K46L, we determined a crystal structure of AM S45R/K46L/S48G/Q50W bound to a maltose binding protein (MBP)-RAMP2-CLR ECD fusion at 1.83 Å resolution (Fig. 3a; Supplementary Table 2) and compared it to our prior structure with wild-type AM (Fig. 3b). Excellent electron density was observed for the AM variant (Supplementary Fig. 1a), however, crystal packing complicated interpretation of the effects of the S45R and K46L substitutions. R356 from MBP stacked on AM K46L and formed an H-bond/ionic bond with RAMP2 E101 resulting in two alternate E101 conformations (Fig. 3a and Supplementary Fig 1b). In addition, packing of a symmetry mate MBP against the α1-β1 loop in CLR caused shifts in the CLR α1 helix and the β1-β2 loop that forms part of the pocket bordered by the RAMP (Supplementary Fig 1c, d). Shifting of RAMP2 enabled E105 to form a salt-bridge with CLR R119 that was not observed in our prior structure (Fig. 3a,b and Supplementary Fig 1c). Electron density for the AM variant S45R side chain supported modeling two alternate conformations, neither of which contacted RAMP2 (Fig. 3a), possibly because of packing effects. To test if S45R forms an ionic interaction with RAMP2 E105 we assessed the ability of S45R to enhance affinity in a cell-based functional cAMP antagonism assay using full-length receptor with RAMP2 E105A. In this assay, the S45R/K46L/S48G/Q50W variant retained ∼10-fold higher affinity than the K46L/S48G/Q50W variant (Fig. 3c). Thus, RAMP2 E105 does not mediate the AM S45R affinity-enhancing effect.

**Figure 3.**
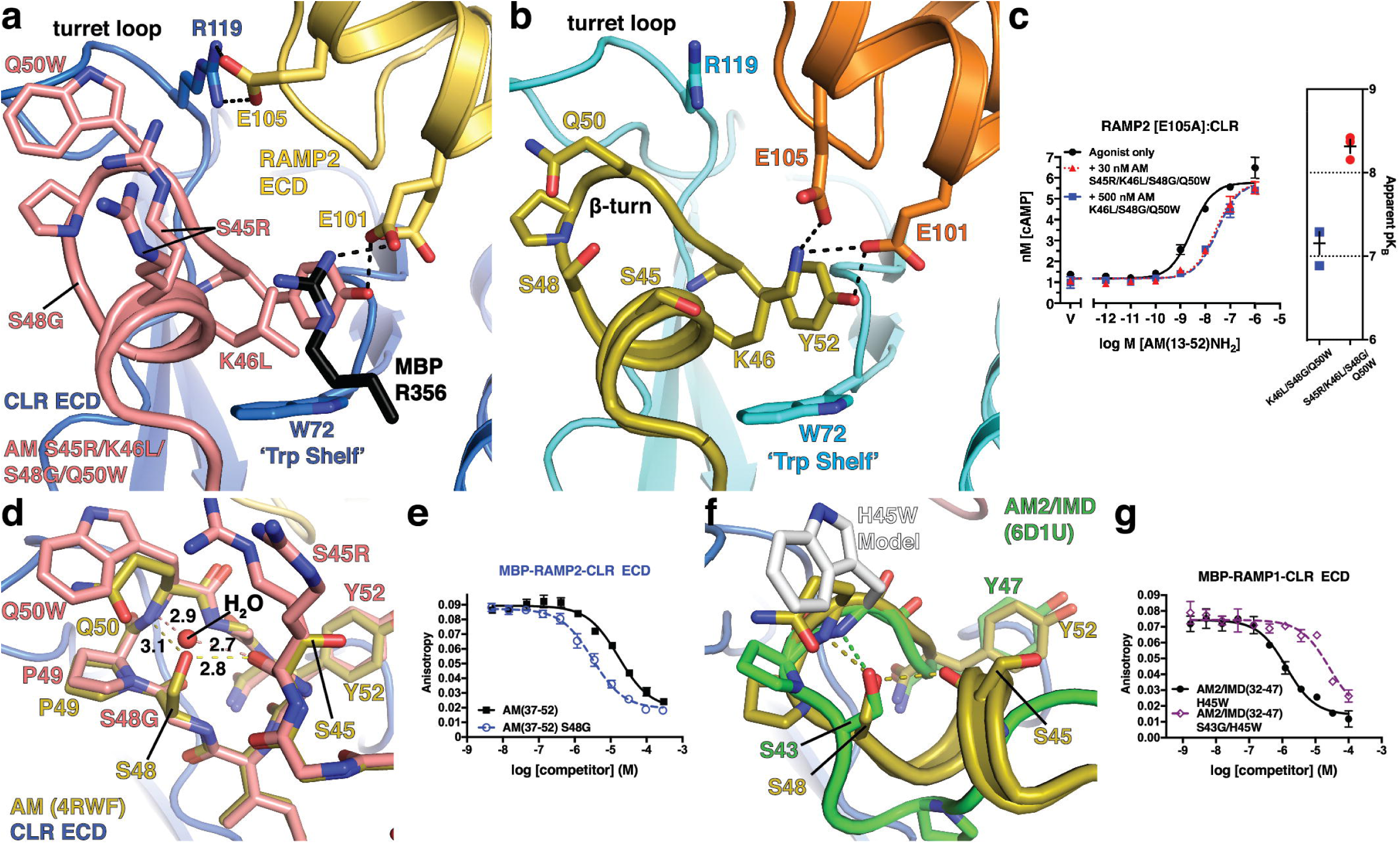
Structural basis for enhanced RAMP2-CLR ECD affinity of AM(37-52) S45R/K46L/S48G/Q50W. (a) 1.83 Å resolution crystal structure of the variant bound to MBP-RAMP2-CLR ECD. (b) Crystal structure of AM-bound MBP-RAMP2-CLR ECD [PDB 4RWF] in the same view as panel (a) for comparison. (c) cAMP signalling antagonism assay in COS-7 cells using RAMP2 E105A:CLR and the indicated concentration of antagonist peptide variants. Right is a scatter plot of mean apparent pK_B_ values determined from three independent experiments with error shown as S.E.M. (d) Superimposition of the new variant structure and 4RWF showing a detailed view of the peptide β-turn. H-bond distances are shown in angstroms. (e) Competition FP assay for the indicated AM peptides at MBP-RAMP2-CLR ECD. (f) Structural alignment of AM [PDB 4RWF] and AM2/IMD [PDB 6D1U]. (g) Competition FP assay for the indicated AM2/IMD peptides at MBP-RAMP1-CLR ECD. Mean pK_I_ ± S.E.M values for the H45W and S43G/H45W variants were 6.08 ± 0.06 and 4.75 ± 0.08, respectively.

Crystal packing did not affect the S48G and Q50W substitutions. Q50W contacted the CLR turret loop and intramolecularly packed against P49, presumably stabilizing the β-turn (Fig. 3d). In the variant structure with S48G, a water molecule occupied the position of the S48 hydroxyl where it formed bridging H-bonds with the main chains of Q50W and S45R similar to the S48 side chain, but with shorter bond distances (Fig. 3d). Flexibility enabled by S48G allowed tighter packing of P49 with Q50W. Supporting this, S48G alone enhanced affinity only ∼5-fold at RAMP2-CLR ECD (Fig. 3e and Supplementary Table 1), rather than the 13-fold enhancement observed with Q50W. Notably, AM2/IMD contains S43 at the position equivalent to AM S48 and we previously showed that the AM2/IMD H45W substitution at the position equivalent to AM Q50 increased affinity (Fig. 3f)^30^. In contrast to our results with AM, however, the S43G substitution in AM2/IMD H45W decreased its ECD affinity ∼20-fold (Fig. 3g).

### Probing the ligand selectivity mechanism by AM K46 and CGRP F37 mutagenesis

The role of AM K46 in selectivity has been difficult to define because it has several structural functions. The aliphatic portion of the side chain intramolecularly packs against Y52 and intermolecularly contacts CLR W72, while the side chain amino group is within H-bond/ionic bond distance of RAMP2 E101 and E105 (Fig. 4a). In mutagenesis studies RAMP2 E101A decreased AM cAMP signaling potency 25-fold, but surprisingly E105A had no effect^31^. It was suggested that the K46 aliphatic contacts were most important and that K46L alteration of selectivity resulted from steric effects rather than lost RAMP2 contact(s)^33^. K46L was thought to push the position 52 side chain into a position like CGRP F37 to improve shape complementarity with the pocket in the RAMP1 complex while decreasing complementarity with the RAMP2 complex pocket (Fig. 4a). The AM library position 46 findings presented thus far and crystal packing limitations on interpretation of K46L shape complementarity effects in the new structure called for revisiting the roles of AM K46 and RAMP allostery in ligand selectivity.

**Figure 4.**
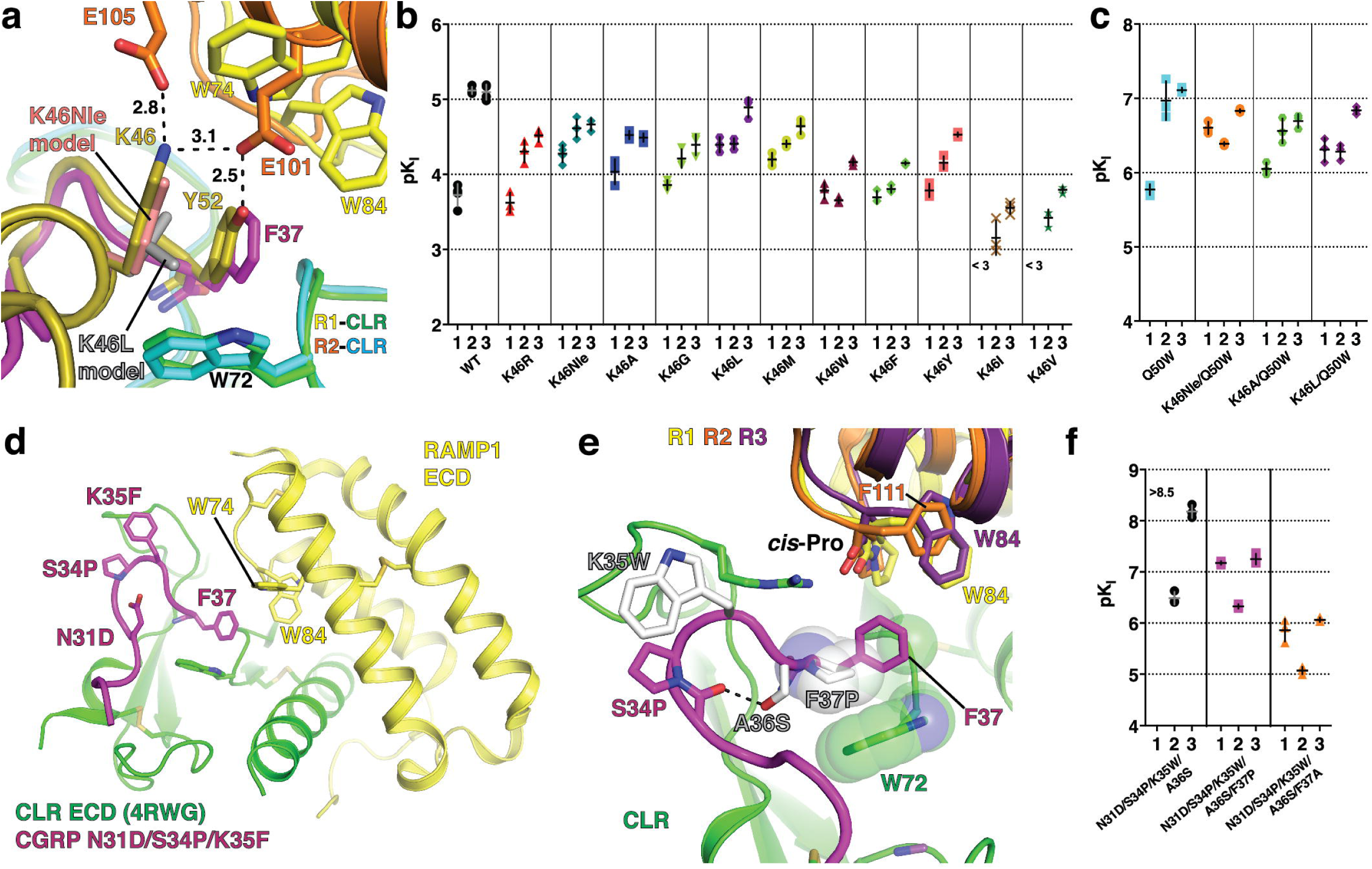
Probing the ligand selectivity mechanism by AM K46 and CGRP F37 mutagenesis. (a) Detailed view of the pocket over the W72 Trp shelf in superimposed AM-bound RAMP2-CLR ECD [4RWF] and CGRP N31D/S34P/K35F-bound RAMP1-CLR ECD [4RWG] crystal structures. AM position 46 substitutions K46Nle and K46L are modeled and key residues are shown as sticks. (b,c) Scatter plots of mean pK_I_ values from competition FP assays with purified MBP-RAMP-CLR ECD complexes and the indicated AM(37-52) K46 variants in the (b) wild-type or (c) Q50W backgrounds. (d) Crystal structure of CGRP N31D/S34P/K35F-bound MBP-RAMP1-CLR ECD [PDB 4RWG] with MBP omitted. (e) CGRP K35W, A36S, and F37P substitutions are modeled and shown with the indicated RAMP1-3 residues that augment the pocket. RAMP3 is a homology model. (f) Scatter plot of mean pK_I_ values from competition FP assays using purified MBP-RAMP-CLR ECD complexes with the indicated CGRP F37 variants in the N31D/S34P/K35W/A36S background. The high-affinity detection limit of the assay prevented unambiguous determination of the affinity of the N31D/S34P/K35W/A36S variant at the RAMP1 complex. See Supplementary Table 3 for the pK_I_ values with SEM, selectivity comparisons, and associated statistical analyses.

AM K46 function was probed by determining the ECD complex affinities of AM(37-52) peptides containing diverse substitutions at position 46 in an otherwise wild-type background (Fig. 4b and Supplementary Table 3). K46R selectively decreased binding with RAMP2/3. The nonstandard amino acid norleucine (Nle) was used to remove the amino group while maintaining aliphatic packing ability (Fig. 4a). K46Nle diminished binding with RAMP2/3 and increased binding with RAMP1 (Fig. 4b). K46A, which further removes some of the intramolecular packing against Y52, was similar to K46Nle, but with slightly decreased affinity at all three complexes. K46G, which removes all side chain intra- and intermolecular contacts had little effect with RAMP1, but decreased binding at RAMP2/3. K46L and K46M increased binding with RAMP1 and decreased binding with RAMP2 while having little or no effect with RAMP3. W, F, and Y selectively decreased binding with RAMP2/3, and I and V decreased binding at all three complexes. K46Nle, K46A, and K46L behaved similarly in the Q50W background (Fig. 4c). These results revealed a crucial role for K46 in selectivity and indicated that its side chain amino group is important for this function.

To examine RAMP allosteric effects we turned to CGRP because it has only a single residue, F37, that contacts the RAMP subunit (Fig. 4d). We reasoned that CGRP peptides that cannot contact the RAMPs would likely still exhibit different ECD complex affinities if the RAMPs stabilize different CLR conformations. Starting with our prior rational design variant CGRP(27-37) N31D/S34P/K35W/A36S that has greatly increased ECD complex affinity^33^, we added the F37P or F37A substitutions to remove contact with the conserved cis-Pro in RAMP1-3 and W84 unique to RAMP1/3 (Fig. 4e). Both variants strongly preferred the RAMP1/3 complexes and the F37P version had higher affinities, presumably due to Pro packing against the Trp shelf (Fig. 4e,f, Supplementary Table 3). These results suggested that RAMP2 stabilized a different CLR ECD conformation than RAMP1/3.

### Antagonism of cAMP signaling by truncated single site ECD-binding and dual site ECD/TMD-binding AM and CGRP variants with enhanced ECD affinities

Using a COS-7 cell-based cAMP signaling assay with transiently expressed RAMP:CLR complexes, we examined the enhanced affinity AM and CGRP variants in the context of truncated antagonist scaffolds that lack the N-terminal disulfide-linked loop required for receptor activation (Fig. 5a,b). In the short ECD-binding fragments we characterized the library-identified AM variants S48G/Q50W, S45R/K46L/Q50W, K46L/S48G/Q50W, and S45R/K46L/S48G/Q50W and the new rationally designed CGRP variant N31D/S34P/K35W/A36S/F37P. Representative assays for AM(37-52) S48G/Q50W are shown in Fig. 5c, and the receptor affinities derived from the functional data are summarized in Fig. 5f, g and Supplementary Table 4. The AM and CGRP ECD-binding antagonist variants exhibited surmountable antagonism with nanomolar apparent K_B_ values that were comparable to their K_I_ values for ECD complex binding.

**Figure 5.**
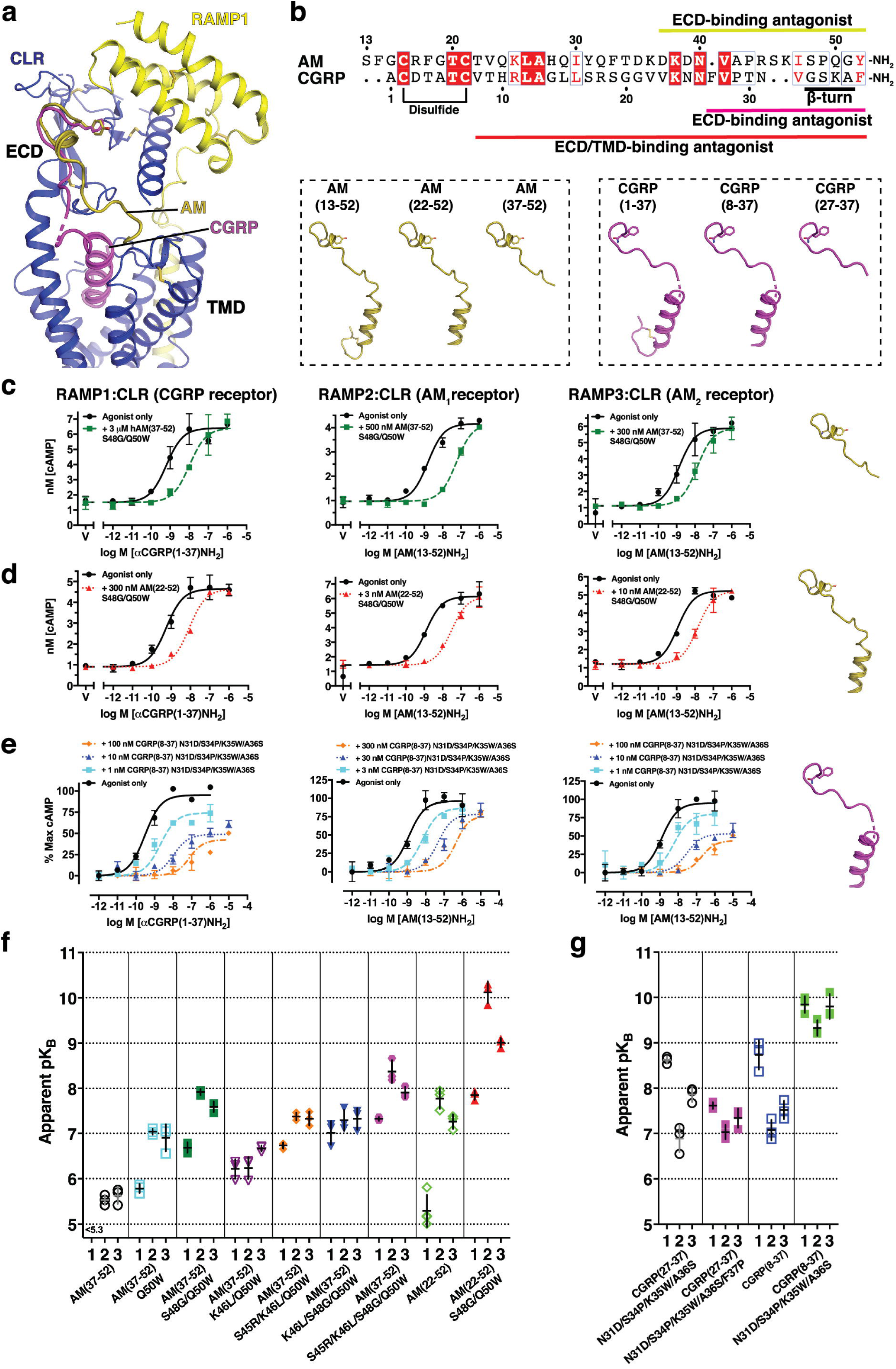
Antagonism of cAMP signaling by single site ECD and dual site ECD/TMD-binding truncated AM and CGRP variants. (a) Cryo-EM structure of CGRP-bound full-length RAMP1:CLR [PDB 6E3Y] with the superimposed AM ECD-binding fragment from 4RWF. (b) Amino acid sequence alignment of AM and CGRP and cartoon depictions of their agonist and N-terminally truncated ECD/TMD-binding and ECD-binding antagonist forms. (c-e) Representative cAMP signaling antagonism assays for (c) AM(37-52) S48G/Q50W, (d) AM(22-52) S48G/Q50W, or (e) CGRP(8-37) N31D/S34P/K35W/A36S at transiently expressed RAMP:CLR complexes in COS-7 cells. (f,g) Scatter plots of mean apparent pK_B_ values for the indicated (f) AM or (g) CGRP antagonist peptides. Values for the peptides with open symbols are from Booe et al, 2018 for comparison^33^. See Supplementary Table 4 for the pK_Bapp_ values with SEM, selectivity comparisons, and associated statistical analyses and Supplementary Table 5 for the hemi-equilibrium model parameters derived from fitting the CGRP(8-37) N31D/S34P/K35W/A36S data.

In the dual site ECD/TMD-binding fragments we tested the AM S48G/Q50W and CGRP N31D/S34P/K35W/A36S variants because these had the highest ECD affinities while retaining selectivities comparable to the wild-type peptides. AM(22-52) S48G/Q50W exhibited surmountable antagonism at all three receptors and a striking 80 pM apparent K_B_ for the AM_1_ receptor while having 170-fold and 12-fold lower affinities at the CGRP and AM_2_ receptors, respectively (Fig. 5d,f and Supplementary Table 4). CGRP(8-37) N31D/S34P/K35W/A36S antagonism was insurmountable at all three receptors (Fig. 5e). In a simultaneous antagonist/agonist addition format rather than antagonist pre-incubation format, little or no depression of the E_max_ was observed (Supplementary Fig. 2), consistent with the insurmountable behavior reflecting hemi-equilibrium due to a slow offset competitive antagonist^25^. Fitting the data to a hemi-equilibrium operational model^36^ yielded estimated K_B_ values of 141, 468, and 158 pM for the RAMP1, −2, and −3 complexes, respectively (Fig. 5g and Supplementary Tables 4 and 5). Very slow off rates of 0.0014, 0.0036, and 0.0018 min^-1^ were estimated that corresponded to residence times of 12.5, 5.2, and 11 hr at the CGRP, AM_1_, and AM_2_ receptors, respectively (Supplementary Table 5).

### AM and CGRP agonist variants with dramatically enhanced ECD affinities exhibit long-acting, sustained cAMP signaling in model cell lines and primary cells

Next, we characterized the AM(13-52) S48G/Q50W and CGRP(1-37) N31D/S34P/K35W/A36S agonist variants in cAMP accumulation assays using COS-7 cells transiently expressing the receptors. Both variants had significantly enhanced potencies at their non-cognate receptors, but showed only minor potency increases at their cognate receptors (Fig. 6a,b and Supplementary Table 6). We reasoned that slow kinetics at the cognate receptors prevented equilibrium from being reached, so we turned to an alternative assay format to measure sustained cAMP signaling after exposure of the cells to 100 nM agonist followed by ligand washout (see Methods). In this format, AM(13-52) S48G/Q50W and CGRP(1-37) N31D/S34P/K35W/A36S yielded significantly enhanced sustained signaling at their cognate AM_1_ and CGRP receptors in COS-7 cells, respectively (Fig. 6c,d). To lesser extents, both variants also had enhanced sustained signaling at the AM_2_ receptor (Supplementary Fig. 3).

**Figure 6.**
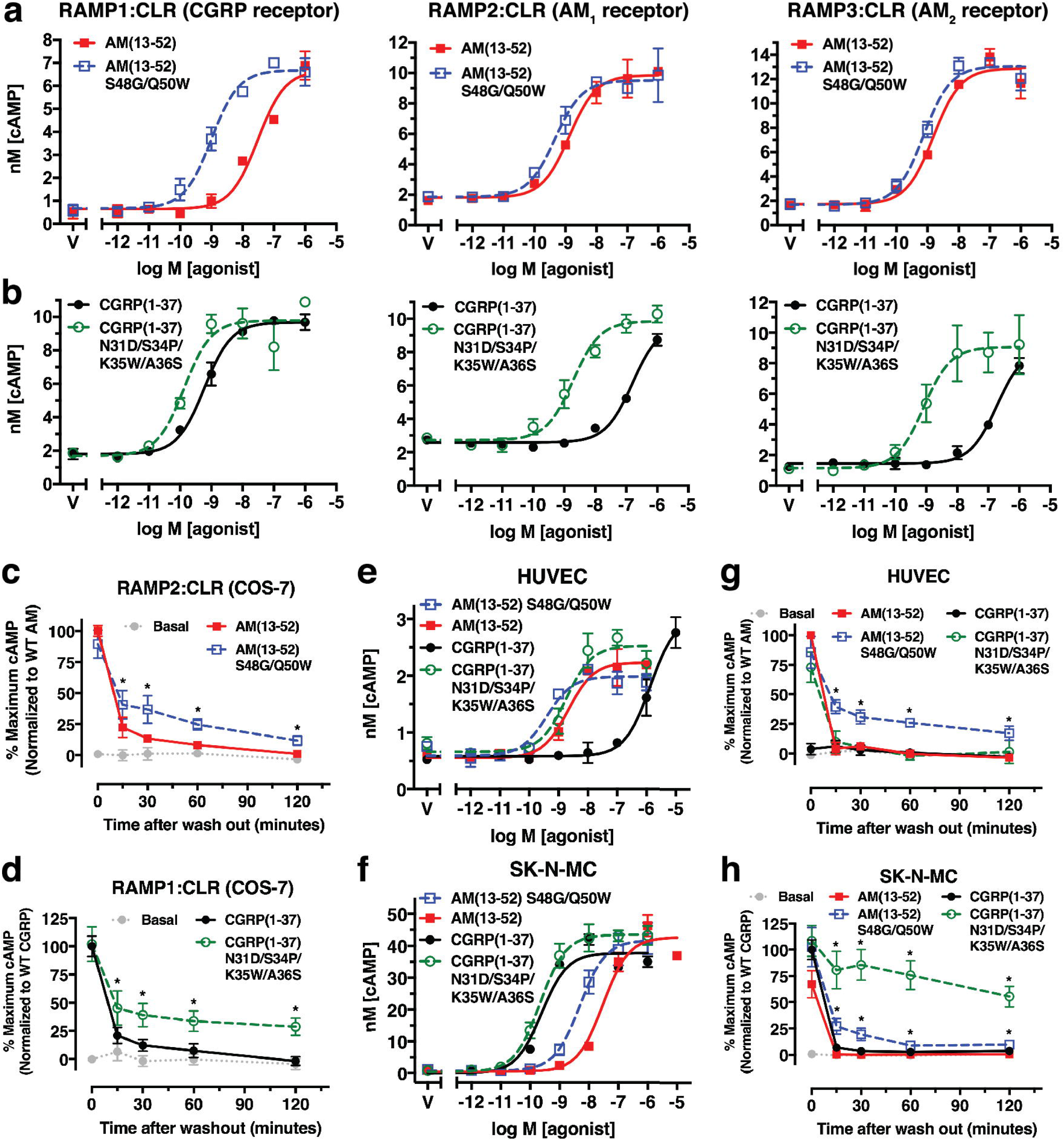
cAMP signaling properties of AM and CGRP agonist variants with enhanced ECD affinities. (a,b) Concentration-response cAMP accumulation assays in COS-7 cells transiently expressing the indicated receptors with the indicated (a) AM or (b) CGRP peptides. (c,d) Sustained cAMP signaling after ligand washout assays using 100 nM of the indicated (c) AM or (d) CGRP peptides at the indicated receptors transiently expressed in COS-7 cells. (e,f) Concentration-response cAMP accumulation assays for the indicated AM and CGRP peptides in (e) primary HUVECs or (f) SK-N-MC cells. (g,h) Sustained cAMP signaling after ligand washout assays with 100 nM of the indicated AM and CGRP peptides in (g) primary HUVECs or (h) SK-N-MC cells. Representative accumulation assays are shown whereas the washout assays are presented as a normalized composite of three independent experiments. See Supplementary Table 6 for the pEC_50_ values with SEM from the accumulation assays and associated statistical analyses.

Last, we characterized the agonist variants using primary human umbilical vein endothelial cells (HUVECs) that express the AM_1_ receptor^37^ and the SK-N-MC neuroblastoma cell line that expresses the CGRP receptor^38^. In standard cAMP accumulation assays the AM and CGRP agonist variants had only slightly enhanced potencies at their cognate receptors in HUVECs or SK-N-MC cells, respectively, and exhibited significantly enhanced potencies at their non-cognate receptors (Fig. 6e, f), similar to the results in COS-7 cells. In washout assays with HUVECs, the wild-type agonists and the CGRP variant failed to elicit sustained cAMP signaling, whereas the AM variant exhibited a significant sustained response with a cAMP level 2 hr post-washout equivalent to ∼25% of the pre-washout level for wild-type AM (Fig. 6g). In SK-N-MC cells the wild-type agonists did not exhibit sustained signaling, the AM variant had a low level of sustained signaling, and the CGRP variant elicited a dramatic sustained response with a cAMP level 2 hr post-washout equivalent to ∼50% of the pre-washout level for wild-type CGRP (Fig. 6h). We term these long-acting, **<u>s</u>**ustained **<u>s</u>**ignaling AM(13-52) S48G/Q50W and CGRP(1-37) N31D/S34P/K35W/A36S agonist variants “ss-AM” and “ss-CGRP”.

## Discussion

Using a powerful combinatorial peptide library approach, novel AM variants with dramatically increased ECD affinities were identified. This work is a substantial advance over our prior rational design effort that identified affinity-enhancing substitutions in the short AM and CGRP ECD-binding fragments^33^. The highest affinity AM variants obtained in that study were relatively non-selective or had altered preference for the CGRP receptor. The AM S48G/Q50W variant identified here exhibited ∼1000-fold increased ECD affinity while maintaining good selectivity for the AM receptors. Notably, we would never have arrived at the S48G substitution by rational design because the S48 side chain makes H-bonds that appeared to be critical for stabilizing the β-turn and α-helical turn (Fig. 3d). The new crystal structure surprisingly revealed that a bridging water is actually better at performing this function (Fig. 3d). That this contributed to the S48G-enhanced affinity is supported by the failure of the equivalent S43G substitution to enhance affinity in AM2/IMD, which lacks the α-helical turn (Fig. 3f). S48G also synergized with Q50W by allowing its tighter intramolecular packing with P49 to stabilize the β-turn. Much of the increased affinity from S48G and Q50W likely results from turn stabilization and the accompanying reduction of the entropic penalty associated with peptide ordering upon receptor binding.

Another novel library-identified substitution, S45R, only enhanced affinity in the defined AM variants when combined with K46L. Crystal packing in the AM S45R/K46L/S48G/Q50W-bound structure prevented us from unambiguously determining how S45R increased affinity, but given that RAMP2 E105 was not involved (Fig. 3c), we speculate that S45R sits over K46L similar to MBP R356 and makes ionic interactions with RAMP2 E101 (Fig. 3a) or RAMP3 E74 (Fig. 1c). This would explain why S45R had no effect in the presence of the wild-type K46 residue, which would block such an interaction. S45R also enhanced affinity at RAMP1-CLR ECD for reasons that are unclear, but the affinity enhancement was largest with RAMP2 and the library screen suggested that S45R favors RAMP2/3. There may be other S45R-containing variants in the library that retain better selectivity for the AM receptors, perhaps by including other K46 substitutions and/or I47L/M, which we did not pursue here.

Significantly, in addition to identifying affinity-enhancing substitutions, the library screen also revealed an important role for AM K46 in receptor selectivity. We previously argued that the main role of K46 is packing against Y52^33^, but the new data do not support this. A variety of small polar or nonpolar residues at position 46 increased binding with RAMP1 and the AM K46 mutagenesis in defined variants highlighted the importance of the K46 amino group for discriminating RAMP2/3 from RAMP1 (Fig. 1d and Fig. 4b,c). AM K46 has a positive role to contact RAMP2 E101 or RAMP3 E74, and it appears to have a negative role at the RAMP1 complex because K46Nle and K46A improved binding with RAMP1. Electrostatics may explain this because RAMP2/3 have more favorable charge complementarity for K46 than RAMP1 (Supplementary Fig. 4d-g) and this is consistent with RAMP mutagenesis data^39^. Thus, AM K46 contacts with RAMP2 E101 and RAMP3 E74 are important for ligand selectivity in addition to the RAMP1 W84, RAMP2 E101, and RAMP3 E74/W84 contacts with the peptide C-terminal residues previously identified^30, 31, 33^. The selectivity-determining RAMP contacts for AM and CGRP are summarized in Supplementary Fig. 4a-c.

The small number of RAMP-peptide contacts in the structures suggested that RAMPs may also have an allosteric role in selectivity^30–32^, but evidence for this has been limited. We leveraged our high-affinity CGRP(27-37) N31D/S34P/K35W/A36S variant^33^ to probe for RAMP-dependent conformational differences in the CLR ECD. The new F37P- or F37A-containing versions of this variant that lacked the ability to contact the RAMP subunits nonetheless preferred RAMP1/3 in the ECD complex binding (Fig. 4f) and full-length receptor antagonism (Fig. 5g) assays. These data suggested that RAMP1/3 stabilize different CLR ECD conformations than RAMP2. This is consistent with the RAMP1- and RAMP2-CLR ECD crystal structures where subtle movement of the CLR β1-β2 loop (Fig. 4a) and more dramatic movement of the CLR R119 side chain (Supplementary Fig. 4a,b) resulted in altered pocket shapes that appeared to be RAMP-dependent^31^. The ECD complex binding data seem to suggest that RAMP3 and RAMP1 stabilize similar CLR ECD conformations (Fig. 4f), but a crystal structure of the RAMP3-CLR ECD complex is needed to verify this. The data now available supports a model in which RAMPs modulate CLR ligand selectivity at the level of the ECD complexes through both direct RAMP-peptide contacts and allosteric modulation of CLR.

Using cAMP signaling assays we characterized key library-identified AM and rationally designed CGRP variants in the context of single site ECD-binding and dual site ECD/TMD-binding competitive antagonist peptide scaffolds (Fig. 5). The single site AM(37-52) and CGRP(27-37) antagonist variants exhibited affinities for the full-length receptors (Fig. 5f, g) that were satisfyingly in agreement with their affinities for the purified ECD complexes (Fig. 2a,b and Fig. 4f). Several of these had nanomolar affinities. Strikingly, the dual site AM(22-52) S48G/Q50W antagonist exhibited picomolar affinity for the AM_1_ receptor and excellent selectivity over the CGRP receptor. CGRP(8-37) N31D/S34P/K35W/A36S was a picomolar affinity, slow offset antagonist that had impressively long estimated receptor residence times, particularly at the CGRP receptor (12.5 hrs). Many of these novel antagonists will be valuable pharmacological tools and some may hold promise as long residence-time therapeutics. Future research is needed to directly measure their binding kinetics and test their activity in appropriate animal models.

Most significantly, incorporating the ECD affinity-enhancing substitutions into AM and CGRP agonist peptide scaffolds generated the sustained signaling agonists ss-AM and ss-CGRP (Fig. 6). In standard cAMP signaling assays they appeared to be non-selective (Fig. 6a,b), however, the washout assays in HUVECs and SK-N-MC cells revealed that they selectively promoted sustained cAMP signaling at their cognate receptors (Fig. 6g,h). The weak sustained signaling observed with ss-AM in SK-N-MC cells may result from low-level AM_1_ expression because RAMP2 mRNA has been detected in this cell line^38, 40^. Going forward it will be important to determine if the sustained signaling results from long receptor residence time at the cell surface, signaling from internalized receptors, or another mechanism^26^. Interestingly, CGRP receptor signaling in pain transmission through PKC and ERK occurred from endosomes^41^, and sustained cAMP signaling from endosomes is a well documented phenomenon with another class B receptor, the parathyroid hormone receptor^42^. Regardless of the mechanism, ss-AM and ss-CGRP have the potential to overcome the short plasma half-lives of the wild-type agonists *in vivo*. Encouragingly, an acylated CGRP analog with prolonged circulatory half-life exhibited cardioprotective effects in mouse models^43^. ss-AM and ss-CGRP hold promise as novel therapeutics with long-acting properties based on long receptor residence times rather than prolonged circulatory half-lives.

## Methods

### Cell culture

COS-7 (CRL 1651), SK-N-MC (HTB-10), and primary human umbilical vein endothelial cells (HUVECs) (PCS-100-010) were from American Type Culture Collection (Manassas, VA, USA). Dulbecco’s modified eagle media (DMEM) with 4.5 g/L glucose and L-glutamine was from Lonza (Basel, Switzerland). Minimal Essential Media (MEM) was from Thermo Fisher Scientific. Vascular Cell Basal Media (PCS-100-030) and Endothelial Cell Growth Kit-BBE (PCS-100-040) were from American Type Culture Collection (Manassas, VA, USA). Fetal bovine serum was from Life Technologies (Carlsbad, CA). COS-7 cells were cultured in DMEM supplemented with 10% fetal bovine serum. SK-N-MC cells were cultured in MEM supplemented with 10% fetal bovine serum. HUVECs were cultured in Vascular Cell Basal Media supplemented with Endothelial Cell Growth Kit-BBE according to manufacturer’s instructions (American Type Culture Collection). All cells were grown at 37°C, 5% CO_2_ in a humidified CO_2_ incubator.

### Plasmids

The mammalian expression plasmids for the tethered ECD fusion proteins maltose binding protein (MBP)-RAMP1.24-111-(GS)_5_-CLR.29-144-H_6_, MBP-RAMP2.55-140[L106R]-(GS)_5_-CLR.29-144-H_6_, and MBP-RAMP3.25-111-(GS)_5_-CLR.29.-144-H_6_, and the bacterial expression plasmid for MBP-RAMP2.55-140[L106R]-(GSA)_3_-CLR.29-144-H_6_ were previously described^30, 31^. Mammalian expression plasmids encoding full-length RAMP1, RAMP2, RAMP3, CLR, FLAG-RAMP2 [E105A], and HA-CLR have been described elsewhere^30, 31^. All plasmids used the human RAMP and CLR sequences.

### Synthetic peptides

The AM(37-52) Q50W positional scanning-synthetic peptide combinatorial library was custom synthesized by RS Synthesis (Louisville, KY, USA). The amino acid sequences for the five positional libraries were: 1) DKDNVAPROXXXPWGX-NH_2_, 2) DKDNVAPRXOXXPWGX-NH_2_, 3) DKDNVAPRXXOXPWGX-NH_2_, 4) DKDNVAPRXXXOPWGX-NH_2_, and 5) DKDNVAPRXXXXPWGO-NH_2_, where O is the defined position with one of 19 natural amino acids (no cysteine) and X is a variable position containing these 19 amino acids in an equimolar mixture. Each positional library comprised 19 distinct mixtures that each contained 130,321 theoretical peptides (19^4^). There were 2,476,099 theoretical peptides total in the library (130,321*19). Each of the 95 crude lyophilized peptide mixtures was reconstituted at 10 mg/mL in 10 % (v/v) DMSO. Peptide mixture concentrations were determined by UV absorbance at 280 nm using molar absorptivity calculated based on Trp and Tyr residues at defined positions, ignoring the contribution from Trp or Tyr residues at the randomized positions.

Defined AM, CGRP, and AM2/IMD variant peptides were custom synthesized and HPLC purified by RS Synthesis (Louisville, KY, USA). Peptides containing the non-standard amino acid norleucine (Nle) were custom synthesized and HPLC purified by New England Peptide (Gardner, MA, USA). Wild-type agonist αCGRP(1-37) and AM(13-52) were from Bachem (Bubendorf, Switzerland). The lyophilized powders were reconstituted at 10 mg/ml in sterile ultrapure water. Peptide concentrations were determined by UV absorbance at 280 nm using the molar absorptivity calculated based on Tyr, Trp, and cystine residues. The concentration of the FITC-(Ahx)-AM(37-52) S45W/Q50W peptide was determined as previously described^33^. If a peptide lacked Tyr or Trp residues, its concentration was determined by assuming 80% peptide content. Peptides were stored as aliquots at −80°C. Supplementary Table 7 lists the sequences of all defined peptides used in this study.

### Purified proteins

The MBP-RAMP1.24-111-(GS)_5_-CLR.29-144-H_6_, MBP-RAMP2.55-140[L106R]-(GS)_5_-CLR.29-144-H_6_, and MBP-RAMP3.25-111-(GS)_5_-CLR.29-144-H_6_ tethered ECD fusion proteins for FP binding assays were expressed in HEK293T cells and purified as described^30^. The MBP-RAMP2.55-140[L106R]-(GSA)_3_-CLR.29-144-H_6_ fusion protein for crystallization was expressed in *E. coli* and purified as previously described^31^. MBP facilitates crystallization of the fusion protein. Purified proteins were stored at −80°C and their concentrations were determined by Bradford assay with a BSA standard curve.

### FP peptide-binding assay

Fluorescence polarization/anisotropy (FP) peptide binding assays using FITC-labeled AM(37-52) S45W/Q50W probe and the three purified, HEK293 cell-produced MBP-RAMP-CLR ECD fusion proteins were performed at room temperature as previously described^30^ except using 100 mM sodium HEPES pH 7.4. For library screening, 10 µL of peptide mixtures at 5X (pre-diluted in reaction buffer) were mixed with 20 µL of a master mix containing the FITC-AM probe followed by addition of 20 µL of a master mix containing purified ECD fusion protein. The final concentration of each unique peptide in the screening assay was 500 pM for RAMP1- and RAMP3-CLR ECD fusion proteins and 1.5 nM for RAMP2-CLR ECD. Raw data were normalized as % of mean of the no competitor control performed in each experiment. The data shown are a composite of normalized data from two independent experiments each performed with duplicate technical replicates.

For defined peptides, competition binding assays were performed using 7 nM FITC-AM probe and 60 nM MBP-RAMP1-CLR ECD, 40 nM MBP-RAMP2-CLR ECD, or 7 nM MBP-RAMP3-CLR ECD and increasing amounts of unlabeled competitor peptides. Equilibrium dissociation constants (K_D_) of the probe for the fusion proteins expressed in HEK293T cells were previously reported^30^. Equilibrium dissociation constants of the unlabeled peptides (K_I_) were determined using nonlinear regression curve fitting to user-defined exact analytical equations expressed in terms of the total ligand and receptor concentrations in Prism v. 7.0d as previously described^30^. For low affinity peptides where the entire competition curve could not be defined, the bottom was constrained to be the same as that of a high-affinity peptide assayed within the same experiment. A Polarstar Omega plate reader was used for the FP measurements (BMG Labtech, Germany).

### Crystallization, diffraction data collection, structure solution, and refinement

Purified, bacterially-expressed MBP-RAMP2 ECD-(GSA)_3_-CLR ECD fusion protein was dialyzed to buffer containing 10 mM Tris-HCl pH 7.5, 50 mM NaCl, 1 mM maltose, 1 mM EDTA and mixed with AM(37-52) S45R/K46L/S48G/Q50W in a 1:1.2 protein:peptide molar ratio, incubated on ice for 1 hour, and then concentrated to 30 mg/mL using a 3,000 Da molecular weight cutoff spin concentrator device (Millipore). Crystals were grown by hanging drop vapor diffusion in 20% (w/v) PEG monomethyl ether (MME) 5000, 0.1 M sodium HEPES pH 8.2, 150 mM sodium formate and 3% (v/v) DMSO. Crystals were cryoprotected by overnight dialysis into a solution containing mother liquor supplemented with 12% (w/v) sucrose then flash frozen in liquid nitrogen. Initial crystals were checked for diffraction on a home X-ray source and then high-resolution diffraction data were remotely collected from a single crystal at 100K at LS-CAT 21-ID-G (λ = 0.9786 Å) of the Advanced Photon Source (Argonne, IL). The data were processed using HKL2000 v. 712^44^ and CCP4 v. 7.0.066^45^. The structure was solved by molecular replacement (MR) with Phaser v. 2.8.2^46^ using MBP with maltose removed (PDB: 3C4M) and RAMP2-CLR ECD with MBP and peptide removed (PDB: 4RWF) as search inputs. The MR solution contained one molecule in the asymmetric unit. The MR solution was rigid body refined with REFMAC5 v. 5.8.0238^47^ by treating MBP, RAMP2 ECD, and CLR ECD as separate rigid bodies. The final model was completed by iterative rounds of manual model building using COOT^48^ and TLS restrained refinement in REFMAC5 v. 5.8.0238^47^. The Ramachandran plot had no outliers and 1.6% and 98.4% of residues were in the allowed and preferred regions, respectively.

### Concentration-response cAMP accumulation assays

COS-7 cells were seeded into 96-well plates (Corning, NY, USA) at a cell density of 20,000 cells/well and co-transfected with human RAMP1-, RAMP2-, or RAMP3- and CLR-encoding plasmids using PEI as previously described^33^. These assays were performed at 37 °C. Forty-eight hours after transfection, the cells were pre-incubated in assay buffer consisting of DMEM supplemented with 0.1% (w/v) fatty acid-free bovine serum albumin (BSA) and 1 mM 3-isobutyl-1-methylxanthine (IBMX) for 30 minutes, followed by continuous exposure to the indicated agonist concentration in assay buffer for 15 minutes. For antagonism experiments, fixed concentrations of antagonists were pre-incubated with cells in assay buffer for 30 minutes followed by continuous exposure to the indicated agonist concentration in the presence of the antagonist for 30 minutes, unless otherwise noted.

SK-N-MC cells and HUVECs were seeded into 96-well plates at cell densities of 15,000 cells/well and 10,000 cells/well, respectively. HUVECs were used at passage four or lower. Forty-eight hours after seeding, SK-N-MC cells and HUVECs were pre-incubated for 30 minutes in Krebs-Ringer-HEPES (KRH) buffer (25 mM HEPES pH 7.4, 104 mM NaCl, 5 mM KCl, 2 mM CaCl_2_, 1.2 mM MgSO_4_, and 1.2 mM KH_2_PO_4_) supplemented with 0.1% (w/v) fatty acid-free BSA and 1 mM IBMX. Cells were then stimulated by continuous exposure to the indicated concentrations of agonist peptides in the same buffer for 15 minutes.

In all cases the cells were lysed with 40 µL of 6% (v/v) perchloric acid and neutralized with sodium bicarbonate and sodium HEPES pH 7.4 for a total lysate volume of 91 µL. A LANCE cAMP detection kit (Perkin-Elmer) was used to quantify cAMP in the lysates according to the manufacturer’s directions as previously described^33^. Six µL of lysate was used in the 24 µL total assay volume and the data are presented as nM cAMP in this assay volume. A Polarstar Omega plate reader was used for the LANCE measurements (BMG Labtech, Germany).

Concentration-response data were fit by nonlinear regression in Prism v. 7.0d (GraphPad) using the log(agonist) vs. response model with a standard Hill slope of 1 to determine agonist potency, pEC_50_. For antagonism assays where surmountable antagonism was observed, the apparent affinity of the antagonist (pK_Bapp_) was determined using the Gaddum/Schild EC50 shift model with the Hill and Schild slopes constrained to 1 assuming competitive antagonism. For insurmountable antagonism, the data were fit to a user-defined hemi-equilibria operational model as previously described:^36^

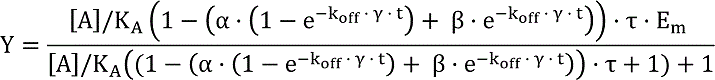

where:

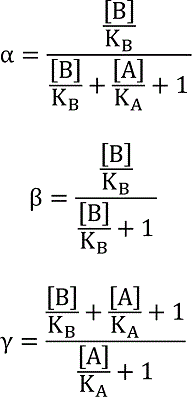

These data were normalized to % maximum cAMP (E_m_) of the agonist curve in the absence of antagonist. Stimulation time (t) was constrained as 30 minutes and the parameters for the equilibrium dissociation constant of the antagonist (K_B_), the equilibrium dissociation constant of the agonist (K_A_), the operational efficacy (τ), and the antagonist dissociation rate (k_off_) were globally fit.

### Sustained cAMP signaling after agonist washout assay

This assay was based on one previously described with modifications^49^. Cells were cultured and seeded into 96-well plates as described above. These assays were performed at room temperature. The cells were washed twice with 100 µL assay buffer (serum-free DMEM + 0.1% fatty acid-free BSA for COS-7 cells or KRH buffer + 0.1% fatty acid-free BSA for HUVECs and SK-N-MC cells), then pre-incubated in assay buffer for 30 minutes followed by stimulation with 100 nM agonist in 100 µL assay buffer for 10 minutes. The agonist-containing buffer was aspirated and the cells were washed with 100 µL of assay buffer three times over a period of ∼3 minutes to wash away unbound agonist and incubated in assay buffer for the indicated times. At the indicated time point the buffer was aspirated and replaced with assay buffer supplemented with 2 mM IBMX for 10 minutes followed by lysate preparation as described above. For the t=0 time point, 100 nM agonist was added to cells in assay buffer containing 2 mM IBMX and the cells were lysed after 10 minutes. The cAMP concentration in the cell lysates was determined as described above. Data were normalized as % maximum cAMP of the wild-type AM agonist (HUVECs or RAMP2-CLR in COS-7) or wild-type CGRP agonist (SK-N-MC or RAMP1-CLR in COS-7) at the t=0 time point with baseline as 0%.

### Statistical analyses

The pK_I_, pK_Bapp_, and pEC_50_ values are reported as means ± S.E.M. from at least three independent experiments performed on separate days. Each independent experiment was performed with duplicate technical replicates. Statistical comparisons of mean pK_I_, pK_Bapp_, or pEC_50_ values were performed using an unpaired student’s *t* test for two groups or one-way ANOVA with Tukey’s or Dunnet’s post-hoc test for three or more groups using Prism v. 7.0d (GraphPad). Selectivity was reported as Δlog of mean values between indicated pairs and error was reported as 95% confidence interval (CI). For the agonist wash-out assays statistical comparisons of wild-type and mutants were performed for each time point using unpaired student’s *t* test. No weighting or outlier detection was used.

## Acknowledgements

The authors thank Drs. Paul Weigel and Paul DeAngelis for suggesting a peptide library approach to identify enhanced affinity variants, Dr. Debbie Hay for helpful suggestions on working with the SK-N-MC cell line, and Dr. Chris Langmead for kindly providing a Prism file for fitting data to the hemi-equilibrium operational model for slow off-rate antagonists and advice on its use. Use of the Advanced Photon Source LS-CAT Sector 21 beamlines was supported by the Michigan Economic Development Corporation and the Michigan Technology Tri-Corridor grant 085P1000817. We thank Zdzislaw Wawrzak for assistance with remote data collection at APS beamline 21-1D-G. Use of the home X-ray source and microscope for crystal imaging at the Biomolecular Structure Core-OKC was supported by an Institutional Development Award (IDeA) from the National Institute of General Medical Sciences of the National Institutes of Health under grant P20GM103640. This work was supported by grants NIH R01GM104251 and OCAST HR16-005 (AAP) and a predoctoral fellowship from the American Heart Association 18PRE33990152 (JMB).

## Author Contributions

JMB and MLW performed experiments. JMB and AAP analyzed data and wrote the manuscript. AAP conceived and managed the project and acquired funding.

