## Supplementary Tables and Figs for "Picomolar affinity antagonist and sustained signaling agonist peptide ligands for the adrenomedullin and calcitonin gene-related peptide receptors"

**Supplementary Table 1. $pK_I$ values for AM variants binding purified RAMP-CLR ECD complexes in the FP assay.**

| AM(37-52)$NH_2$ peptides | MBP-RAMP1-CLR ECD $pK_I$ ± SEM ($K_I$, nM)[a] | MBP-RAMP2-CLR ECD $pK_I$ ± SEM ($K_I$, nM)[a] | MBP-RAMP3-CLR ECD $pK_I$ ± SEM ($K_I$, nM)[a] | Selectivity, R1 vs. R2, Δlog (95% CI)[c] | Selectivity, R1 vs. R3, Δlog (95% CI)[c] | Selectivity, R2 vs. R3, Δlog (95% CI)[c] |
|---|---|---|---|---|---|---|
| **WT**[b] | 3.73 ± 0.07 (186,000) | 5.12 ± 0.02 (7,590) | 5.10 ± 0.05 (7,940) | -1.39 (-1.61 to -1.18)* | -1.37 (-1.58 to -1.16)* | +0.02 (-0.19 to +0.23) |
| **S48G**[b] | 4.41 ± 0.07 (38,900)* | 5.82 ± 0.04 (1,510)* | 5.76 ± 0.05 (1,740)* | -1.41 (-1.64 to -1.18)* | -1.34 (-1.57 to -1.11)* | +0.06 (-0.17 to +0.29) |
| **Q50W** | 5.77 ± 0.04 (1,700) | 6.97 ± 0.15 (107) | 7.11 ± 0.02 (77.6) | -1.20 (-1.59 to -0.80)* | -1.34 (-1.74 to -0.94)* | -0.14 (-0.54 to +0.25) |
| **S45W/Q50W** | 7.04 ± 0.07 (91.2)* | 7.19 ± 0.08 (64.6) | 8.17 ± 0.02 (6.76)* | -0.15 (-0.42 to +0.13) | -1.12 (-1.40 to -0.85)* | -0.98 (-1.25 to -0.71)* |
| **K46L/Q50W** | 6.31 ± 0.09 (490)* | 6.28 ± 0.06 (525)* | 6.84 ± 0.03 (145)* | +0.03 (-0.26 to +0.31) | -0.53 (-0.82 to -0.24)* | -0.56 (-0.84 to -0.27)* |
| **S45W/K46L/Q50W** | 7.85 ± 0.13 (14.1)* | 7.43 ± 0.12 (34.7)* | 8.21 ± 0.01 (6.17)* | +0.42 (-0.02 to +0.87) | -0.36 (-0.80 to +0.09) | -0.78 (-1.26 to -0.30)* |
| **S45W/K46L/Q50W/Y52F** | 8.20 ± 0.14 (6.31)* | 5.89 ± 0.04 (1,290)* | 7.80 ± 0.01 (15.8)* | +2.31 (+1.89 to +2.74)* | +0.41 (-0.02 to +0.83) | -1.90 (-2.36 to -1.45)* |
| **S45R/Q50W** | 5.88 ± 0.11 (1,320) | 7.44 ± 0.10 (36.3)* | 7.30 ± 0.04 (50.1) | -1.57 (-1.96 to -1.18)* | -1.42 (-1.82 to -1.03)* | +0.15 (-0.25 to +0.54) |
| **S45R/K46L/Q50W** | 7.10 ± 0.03 (79.4)* | 7.48 ± 0.07 (33.1)* | 7.52 ± 0.06 (30.2)* | -0.38 (-0.62 to -0.14)* | -0.42 (-0.66 to -0.18)* | -0.04 (-0.28 to +0.20) |
| **S45R/K46W/Q50W** | 6.38 ± 0.05 (417)* | 6.52 ± 0.02 (302)* | 6.91 ± 0.05 (123) | -0.14 (-0.33 to +0.05) | -0.53 (-0.72 to -0.34)* | -0.39 (-0.58 to -0.20)* |
| **S45W/K46W/Q50W** | 6.93 ± 0.06 (117)* | 6.39 ± 0.06 (407)* | 7.12 ± 0.06 (75.9) | +0.54 (+0.27 to +0.81)* | -0.19 (-0.46 to +0.08) | -0.73 (-1.00 to -0.46)* |
| **S45W/K46G/Q50W/Y52F** | 8.33 ± 0.14 (4.68)* | 6.03 ± 0.03 (933)* | 7.53 ± 0.03 (29.5)* | +2.30 (+1.87 to +2.73)* | +0.80 (+0.37 to +1.23)* | -1.50 (-1.96 to -1.04)* |
| **S48G/Q50W** | 6.62 ± 0.10 (240)* | 8.10 ± 0.03 (7.94)* | 7.86 ± 0.05 (13.8)* | -1.48 (-1.79 to -1.18)* | -1.24 (-1.54 to -0.93)* | +0.24 (-0.06 to +0.55) |
| **S45R/S48G/Q50W** | 6.45 ± 0.04 (355)* | 8.19 ± 0.12 (6.46)* | 7.98 ± 0.03 (10.5)* | -1.74 (-2.05 to -1.42)* | -1.52 (-1.84 to -1.21)* | +0.21 (-0.10 to +0.53) |
| **K46L/S48G/Q50W** | 7.16 ± 0.06 (69.2)* | 7.15 ± 0.02 (70.8) | 7.69 ± 0.10 (20.4)* | +0.01 (-0.29 to +0.29) | -0.53 (-0.82 to -0.24)* | -0.53 (-0.82 to -0.23)* |
| **S45R/K46L/S48G/Q50W** | 7.76 ± 0.07 (17.4)* | 8.35 ± 0.13 (4.47)* | 8.06 ± 0.07 (8.71)* | -0.59 (-1.01 to -0.17)* | -0.30 (-0.72 to +0.12) | +0.29 (-0.13 to +0.71) |
| **S45R/K46A/S48G/Q50W** | 6.55 ± 0.10 (282)* | 7.17 ± 0.02 (67.6) | 7.10 ± 0.05 (79.4) | -0.62 (-0.90 to -0.33)* | -0.55 (-0.84 to -0.27)* | +0.06 (-0.22 to +0.35) |

* $P < 0.05$.

[a]Statistical comparison of $pK_I$ of AM Q50W vs. other mutant AM peptides for each ECD complex was done using one-way ANOVA with Dunnett's post-hoc test.

[b]Statistical comparison of $pK_I$ of AM WT compared to S48G for each ECD complex was using unpaired *t* test.

[c]Statistical comparison of $pK_I$ values between RAMP1-, RAMP2-, and RAMP3-CLR ECDs was performed using one-way ANOVA with Tukey's post-hoc test.

**Supplementary Table 2. X-Ray Data collection and refinement statistics**

| | MBP-RAMP2.55-140[L106R]-(GSA)$_3$-CLR.29-144-H$_6$:hAM(37-52)NH$_2$ S45R/K46L/S48G/Q50W |
|---|---|
| **Data collection** | |
| Space group | $P2_12_12_1$ |
| Cell dimensions | |
| *a*, *b*, *c* (Å) | 68.02, 84.99, 123.06 |
| a, b, g (°) | 90, 90, 90 |
| Resolution (Å) | 50.0 - 1.83 (1.86 – 1.83)* |
| $R_{merge}$ | 0.087 (1.18) |
| $I$ / s$I$ | 22.7 (1.32) |
| Completeness (%) | 99.4 (98.9) |
| Redundancy | 7.4 (6.9) |
| **Refinement** | |
| Resolution (Å) | 36.97 - 1.83 (1.86 – 1.83)* |
| No. reflections | 59,946 |
| $R_{work}$ / $R_{free}$ | 0.1565 / 0.1896 |
| No. atoms | |
| Protein | 4487 |
| Peptide | 143 |
| Ligand/ion | 32 |
| Water | 602 |
| *B*-factors | |
| Protein | 30.7 |
| Peptide | 31.6 |
| Ligand/ion | 36.3 |
| Water | 37.7 |
| R.m.s. deviations | |
| Bond lengths (Å) | 0.0095 |
| Bond angles (°) | 1.5297 |

*Values in parentheses are for highest-resolution shell.

**Supplementary Table 3. $pK_I$ values for rationally designed AM and CGRP variants binding purified RAMP-CLR ECD complexes in the FP assay.**

| $AM(37\text{-}52)NH_2$ peptides | MBP-RAMP1-CLR ECD $pK_I$ ± SEM ($K_I$, nM)[a] | MBP-RAMP2-CLR ECD $pK_I$ ± SEM ($K_I$, nM)[a] | MBP-RAMP3-CLR ECD $pK_I$ ± SEM ($K_I$, nM)[a] | Selectivity, R1 vs. R2, Δlog (95% CI)[b] | Selectivity, R1 vs. R3, Δlog (95% CI)[b] | Selectivity, R2 vs. R3, Δlog (95% CI)[b, c] |
|---|---|---|---|---|---|---|
| **WT** | 3.73 ± 0.07 (186,000) | 5.12 ± 0.02 (7,590) | 5.10 ± 0.05 (7,940) | -1.39 (-1.61 to -1.18)* | -1.37 (-1.58 to -1.16)* | +0.02 (-0.19 to +0.23) |
| **K46R** | 3.62 ± 0.08 (240,000) | 4.30 ± 0.09 (50,100)* | 4.52 ± 0.05 (30,200)* | -0.68 (-1.01 to -0.37)* | -0.90 (-1.22 to -0.58)* | -0.22 (-0.53 to +0.10) |
| **K46Nle** | 4.28 ± 0.06 (52,500)* | 4.62 ± 0.07 (24,000)* | 4.67 ± 0.05 (21,400)* | -0.34 (-0.56 to -0.12)* | -0.39 (-0.63 to -0.15)* | -0.05 (-0.29 to +0.19) |
| **K46A** | 4.04 ± 0.10 (91,200)* | 4.53 ± 0.03 (29,500)* | 4.49 ± 0.03 (32,400)* | -0.49 (-0.76 to -0.22)* | -0.46 (-0.73 to -0.18)* | +0.03 (-0.24 to +0.30) |
| **K46G** | 3.86 ± 0.04 (138,000) | 4.21 ± 0.07 (61,700)* | 4.40 ± 0.08 (39,800)* | -0.35 (-0.64 to -0.07)* | -0.54 (-0.83 to -0.25)* | -0.18 (-0.47 to +0.11) |
| **K46L** | 4.40 ± 0.06 (39,800)* | 4.41 ± 0.04 (38,900)* | 4.89 ± 0.08 (12,900) | -0.01 (-0.28 to +0.26) | -0.50 (-0.77 to -0.23)* | -0.49 (-0.76 to -0.22)* |
| **K46M** | 4.20 ± 0.05 (63,100)* | 4.41 ± 0.03 (38.900)* | 4.64 ± 0.06 (22,900)* | -0.21 (-0.42 to +0.01) | -0.44 (-0.65 to -0.23)* | -0.23 (-0.44 to -0.03)* |
| **K46W** | 3.79 ± 0.06 (162,000) | 3.66 ± 0.02 (219,000)* | 4.16 ± 0.03 (69,200)* | +0.13 (-0.05 to +0.31) | -0.38 (-0.56 to -0.20)* | -0.51 (-0.69 to -0.33)* |
| **K46F** | 3.70 ± 0.05 (200,000) | 3.81 ± 0.03 (155,000)* | 4.15 ± 0.01 (70,800)* | -0.11 (-0.26 to +0.04) | -0.45 (-0.60 to -0.31)* | -0.34 (-0.49 to -0.20)* |
| **K46Y** | 3.79 ± 0.05 (162,000) | 4.15 ± 0.05 (70,800)* | 4.53 ± 0.02 (29,500)* | -0.37 (-0.55 to -0.18)* | -0.74 (-0.93 to -0.56)* | -0.38 (-0.56 to -0.19)* |
| **K46I** | < 3 (>1 mM) | 3.16 ± 0.13 (692,000)* | 3.55 ± 0.05 (282,000)* | R2 | R3 | -0.40 (-0.01 to +0.79)* |
| **K46V** | < 3 (>1 mM) | 3.42 ± 0.06 (380,000)* | 3.80 ± 0.03 (158,000)* | R2 | R3 | -0.38 (-0.18 to -0.57)* |
| **Q50W** | 5.77 ± 0.04 (1,700) | 6.97 ± 0.15 (107) | 7.11 ± 0.02 (77.6) | -1.20 (-1.59 to -0.80)* | -1.34 (-1.74 to -0.94)* | -0.14 (-0.54 to +0.25) |
| **K46Nle/Q50W** | 6.60 ± 0.05 (251)* | 6.39 ± 0.01 (407)* | 6.83 ± 0.01 (148)* | +0.22 (+0.08 to +0.34)* | -0.23 (-0.36 to +0.10)* | -0.45 (-0.57 to -0.31)* |
| **K46A/Q50W** | 6.05 ± 0.05 (891)* | 6.56 ± 0.10 (275)* | 6.70 ± 0.06 (200)* | -0.51 (-0.82 to -0.21)* | -0.65 (-0.95 to -0.34)* | -0.13 (-0.44 to +0.17) |
| **K46L/Q50W** | 6.31 ± 0.09 (490)* | 6.28 ± 0.06 (525)* | 6.84 ± 0.03 (145)* | +0.03 (-0.26 to +0.32) | -0.53 (-0.82 to -0.24)* | -0.56 (-0.85 to -0.27)* |
| **$CGRP(27\text{-}37)NH_2$ peptides** | **MBP-RAMP1-CLR ECD $pK_I$ ± SEM ($K_I$, nM)** | **MBP-RAMP2-CLR ECD $pK_I$ ± SEM ($K_I$, nM)[a]** | **MBP-RAMP3-CLR ECD $pK_I$ ± SEM ($K_I$, nM)[a]** | **Selectivity, R1 vs. R2, Δlog (95% CI)[b]** | **Selectivity, R1 vs. R3, Δlog (95% CI)[b]** | **Selectivity, R2 vs. R3, Δlog (95% CI)[b, c]** |
| **N31D/S34P/K35W/A36S** | >8.5 | 6.49 ± 0.07 (324) | 8.18 ± 0.07 (6.61) | R1 | R1 | -1.69 (-1.41 to -1.98)* |
| **N31D/S34P/K35W/A36S/F37P** | 7.18 ± 0.02 (66.1) | 6.33 ± 0.03 (468) | 7.25 ± 0.06 (56.2)* | +0.85 (+0.68 to +1.02)* | -0.08 (-0.25 to +0.10) | -0.92 (-1.10 to -0.75)* |
| **N31D/S34P/K35W/A36S/F37A** | 5.86 ± 0.13 (1,380) | 5.07 ± 0.04 (8,510)* | 6.06 ± 0.03 (871)* | +0.79 (+0.44 to +1.13)* | -0.20 (-0.55 to +0.14) | -0.99 (-1.34 to -0.65)* |

* $P < 0.05$.

[a]Statistical comparison of $pK_I$ of AM WT vs. single mutants or Q50W vs. double mutants for each ECD complex was done using one-way ANOVA with Dunnett's post-hoc test.

[b]Statistical comparison of $pK_I$ values between RAMP1-, RAMP2-, and RAMP3-CLR ECDs was performed using one-way ANOVA with Tukey's post-hoc test.

[c]Statistical comparison of $pK_I$ values between RAMP2- and RAMP3-CLR ECDs was performed using an unpaired *t* test.

**Supplementary Table 4. Apparent $pK_B$ values for AM and CGRP antagonist variants determined by COS-7 cell-based signaling assay**

| Peptide | RAMP1:CLR (CGRP receptor) $pK_{Bapp}$ ± SEM ($K_B$, nM)[a] | RAMP2:CLR ($AM_1$ receptor) $pK_{Bapp}$ ± SEM ($K_B$, nM)[a] | RAMP3:CLR ($AM_2$ receptor) $pK_{Bapp}$ ± SEM ($K_B$, nM)[a] | Selectivity, R1 vs. R2 Δlog (95% CI)[e] | Selectivity, R2 vs. R3 Δlog (95% CI)[e] | Selectivity, R1 vs. R3 Δlog (95% CI)[e] |
|---|---|---|---|---|---|---|
| AM(37-52)[b] | [c]< 5.3 (>5,000) | 5.53 ± 0.06 (2,950)* | 5.62 ± 0.10 (2,400)* | R2 | -0.09 (-0.25 to +0.43) | R3 |
| Q50W[b] | 5.78 ± 0.06 (1,660) | 7.04 ± 0.04 (91.2) | 6.90 ± 0.17 (126) | -1.26 (-1.72 to -0.79)* | +0.13 (-0.34 to +0.60) | -1.12 (-1.59 to -0.65)* |
| S48G/Q50W[d] | 6.69 ± 0.07 (204)* | 7.92 ± 0.04 (12)* | 7.60 ± 0.07 (25.1)* | -1.23 (-1.49 to -0.98)* | +0.32 (+0.07 to +0.58)* | -0.91 (-1.17 to -0.65)* |
| K46L/Q50W[b] | 6.22 ± 0.12 (603)* | 6.23 ± 0.14 (589)* | 6.67 ± 0.04 (214) | -0.28 (-0.91 to +0.34) | -0.31 (-0.93 to +0.31) | -0.03 (-0.65 to +0.59) |
| S45R/K46L/Q50W | 6.73 ± 0.04 (186)* | 7.38 ± 0.05 (41.7) | 7.33 ± 0.09 (46.8) | -0.65 (-0.92 to -0.37)* | +0.05 (-0.23 to +0.33) | -0.60 (-0.88 to -0.32)* |
| K46L/S48G/Q50W | 7.01 ± 0.15 (97.7)* | 7.30 ± 0.14 (50.1) | 7.32 ± 0.14 (47.9) | -0.28 (-0.91 to +0.34) | +0.03 (-0.65 to +0.59) | -0.31 (-0.93 to +0.31) |
| S45R/K46L/S48G/Q50W | 7.33 ± 0.03 (46.8)* | 8.37 ± 0.15 (4.27)* | 7.91 ± 0.08 (12.3)* | -1.05 (-1.47 to -0.63)* | +0.47 (+0.04 to +0.89)* | -0.58 (-1.0 to -0.16)* |
| AM(22-52)[b, f] | 5.29 ± 0.25 (5,130) | 7.78 ± 0.13 (16.6) | 7.27 ± 0.09 (53.7) | -2.45 (-3.2 to -1.71)* | +0.51 (-0.23 to +1.25) | -1.94 (-2.7 to -1.20)* |
| S48G/Q50W[f] | 7.86 ± 0.06 (13.8)* | 10.1 ± 0.13 (0.0794)* | 9.01 ± 0.06 (0.977)* | -2.28 (-2.7 to -1.9)* | +1.12 (+0.74 to +1.50)* | -1.16 (-1.54 to -0.77)* |
| CGRP(27-37) | | | | | | |
| N31D/S34P/K35W/A36S[b] | 8.62 ± 0.05 (2.4) | 6.89 ± 0.17 (129) | 7.86 ± 0.10 (13.8) | +1.73 (+1.22 to +2.25)* | -0.97 (-1.48 to -0.46)* | +0.76 (+0.25 to +1.27)* |
| N31D/S34P/K35W/A36S/F37P | 7.62 ± 0.04 (24)* | 7.03 ± 0.09 (93.3) | 7.34 ± 0.12 (45.7)* | +0.58 (+0.18 to +0.99)* | +0.28 (-0.13 to +0.68) | -0.31 (-0.71 to +0.09) |
| CGRP(8-37)[b, f] | 8.74 ± 0.19 (1.82) | 7.08 ± 0.13 (83.2) | 7.52 ± 0.12 (30.2) | +1.66 (+1.01 to +2.30)* | -0.44 (-1.09 to +0.20) | +1.21 (+0.57 to +1.86)* |
| N31D/S34P/K35W/A36S[f, g] | 9.85 ± 0.11 (0.141)* | 9.33 ± 0.10 (0.468)* | 9.80 ± 0.16 (0.158)* | +0.52 (-0.02 to +1.06) | -0.48 (-1.02 to +0.07) | +0.04 (-0.50 to +0.58) |

* $P < 0.05$.

[a] Statistical comparison of $pK_{Bapp}$ of AM(37-52) Q50W vs. mutant AM peptides for each receptor was done using one-way ANOVA with Dunnett's post-hoc test.

[b] Data previously reported in Booe et al. (2018). *Mol. Pharmacol*.

[c] < 5.3 indicates less than 2-fold shift at 10 μM antagonist concentration.

[d] Used 15 min stimulation time instead of 30 minutes.

[e] Statistical comparison of $pK_{Bapp}$ values between RAMP1-, RAMP2-, and RAMP3:CLR receptors was performed using one-way ANOVA with Tukey's post-hoc test.

[f] Statistical comparison of $pK_{Bapp}$ of AM(22-52) wildtype vs. mutant or CGRP(8-37) wildtype vs. mutant peptides for each receptor was done using unpaired *t* test.

[g] Calculation of $pK_{Bapp}$ was performed using Hemi-equilibrium model (see Methods)

**Supplementary Table 5. Parameters from hemi-equilibrium operational model fitting the αCGRP(8-37) N31D/S34P/K35W/A36S antagonism of cAMP signaling in COS-7 cells.**

| Parameters | RAMP1:CLR (CGRP receptor) | RAMP2:CLR ($AM_1$ receptor) | RAMP3:CLR ($AM_2$ receptor) |
|---|---|---|---|
| $pK_{Bapp}$ ± SEM ($K_B$, pM) | 9.85 ± 0.11 (141) | 9.33 ± 0.10 (468) | 9.80 ± 0.16 (158) |
| $k_{off}$ ± SEM ($min^{-1}$) | 0.0014 ± 0.00011 | 0.0036 ± 0.00075 | 0.0018 ± 0.00058 |
| $pK_A$ ± SEM | 8.14 ± 0.11 | 7.53 ± 0.13 | 7.56 ± 0.02 |
| Log(tau) ± SEM | 1.25 ± 0.03 | 1.41 ± 0.11 | 1.23 ± 0.05 |
| $k_{on}$ (x $10^7$) ± SEM ($M^{-1}min^{-1}$)[a] | 1.02 ± 0.29 | 0.83 ± 0.28 | 1.65 ± 1.11 |
| Residence time ± SEM (min)[b] | 750 ± 56 | 314 ± 79 | 663 ± 167 |
| Half-life ($t_{1/2}$) ± SEM (min)[c] | 520 ± 39 | 218 ± 55 | 459 ± 115 |

[a] $k_{on}$ calculated as $k_{off}$ / $K_{Bapp}$

[b] Residence time calculated as $1/k_{off}$

[c] Half-life calculated as $0.693/k_{off}$

**Supplementary Table 6. $pEC_{50}$ values of AM and CGRP agonist variants from cell-based cAMP signaling assays**

| | **COS-7 cells** | | | **SK-N-MC $pEC_{50} \pm$ SEM ($EC_{50}$, nM)[a]** | **HUVEC $pEC_{50} \pm$ SEM ($EC_{50}$, nM)[a]** |
|---|---|---|---|---|---|
| | **RAMP1:CLR (CGRP receptor) $pEC_{50} \pm$ SEM ($EC_{50}$, nM)[a]** | **RAMP2:CLR ($AM_1$ receptor) $pEC_{50} \pm$ SEM ($EC_{50}$, nM)[a]** | **RAMP3:CLR ($AM_2$ receptor) $pEC_{50} \pm$ SEM ($EC_{50}$, nM)[a]** | | |
| AM(13-52) | 7.31 ± 0.10 (49) | 8.89 ± 0.13 (1.29) | 8.67 ± 0.07 (2.14) | 7.60 ± 0.04 (25.1) | 8.76 ± 0.09 (1.74) |
| S48G/Q50W | 8.78 ± 0.15 (1.66)* | 9.34 ± 0.11 (0.457) | 9.04 ± 0.06 (0.912)* | 8.42 ± 0.11 (3.8)* | 9.44 ± 0.13 (0.363)* |
| CGRP(1-37) | 9.37 ± 0.07 (0.427) | 6.58 ± 0.15 (263) | 6.63 ± 0.08 (234) | 9.51 ± 0.06 (0.309) | < 6 (>1,000) |
| N31D/S34P/K35W/A36S | 10.04 ± 0.09 (0.0912)* | 8.78 ± 0.05 (1.66)* | 8.88 ± 0.09 (1.32)* | 9.71 ± 0.05 (0.195)* | 8.68 ± 0.08 (2.09) |

* $P < 0.05$.

[a] Statistical comparison of $pEC_{50}$ values of AM WT vs. S48G/Q50W or CGRP WT vs. N31D/S34P/K35W/A36S was done using unpaired *t* test.

**Supplementary Table 7. List of peptide sequences.**

| Peptide name | Peptide sequence* |
|---|---|
| AM(13-52)$NH_2$ | SFGCRFGTCTVQKLAHQIYQFTDKDKDNVAPRSKISPQGY-$NH_2$ |
| AM(13-52)$NH_2$ S48G/Q50W **"ss-AM"** | SFGCRFGTCTVQKLAHQIYQFTDKDKDNVAPRSKIGPWGY-$NH_2$ |
| AM(22-52)$NH_2$ | TVQKLAHQIYQFTDKDKDNVAPRSKISPQGY-$NH_2$ |
| AM(22-52)$NH_2$ S48G/Q50W | TVQKLAHQIYQFTDKDKDNVAPRSKIGPWGY-$NH_2$ |
| AM(37-52)$NH_2$ | DKDNVAPRSKISPQGY-$NH_2$ |
| AM(37-52)$NH_2$ Q50W | DKDNVAPRSKISPWGY-$NH_2$ |
| AM(37-52)$NH_2$ S48G | DKDNVAPRSKIGPQGY-$NH_2$ |
| AM(37-52)$NH_2$ K46R | DKDNVAPRSRISPQGY-$NH_2$ |
| AM(37-52)$NH_2$ K46Nle | DKDNVAPRS(Nle)ISPQGY-$NH_2$ |
| AM(37-52)$NH_2$ K46A | DKDNVAPRSAISPQGY-$NH_2$ |
| AM(37-52)$NH_2$ K46G | DKDNVAPRSGISPQGY-$NH_2$ |
| AM(37-52)$NH_2$ K46L | DKDNVAPRSLISPQGY-$NH_2$ |
| AM(37-52)$NH_2$ K46M | DKDNVAPRSMISPQGY-$NH_2$ |
| AM(37-52)$NH_2$ K46W | DKDNVAPRSWISPQGY-$NH_2$ |
| AM(37-52)$NH_2$ K46F | DKDNVAPRSFISPQGY-$NH_2$ |
| AM(37-52)$NH_2$ K46Y | DKDNVAPRSYISPQGY-$NH_2$ |
| AM(37-52)$NH_2$ K46I | DKDNVAPRSIISPQGY-$NH_2$ |
| AM(37-52)$NH_2$ K46V | DKDNVAPRSVISPQGY-$NH_2$ |
| AM(37-52)$NH_2$ S48G/Q50W | DKDNVAPRSKIGPWGY-$NH_2$ |
| AM(37-52)$NH_2$ K46Nle/Q50W | DKDNVAPRS(Nle)ISPWGY-$NH_2$ |
| AM(37-52)$NH_2$ K46A/Q50W | DKDNVAPRSAISPWGY-$NH_2$ |
| AM(37-52)$NH_2$ K46L/Q50W | DKDNVAPRSLISPWGY-$NH_2$ |
| AM(37-52)$NH_2$ S45W/Q50W | DKDNVAPRWKISPWGY-$NH_2$ |
| AM(37-52)$NH_2$ S45R/Q50W | DKDNVAPRRKISPWGY-$NH_2$ |
| AM(37-52)$NH_2$ S45R/K46L/Q50W | DKDNVAPRRLISPWGY-$NH_2$ |
| AM(37-52)$NH_2$ S45R/S48G/Q50W | DKDNVAPRRKIGPWGY-$NH_2$ |
| AM(37-52)$NH_2$ S45R/K46W/Q50W | DKDNVAPRRWISPWGY-$NH_2$ |
| AM(37-52)$NH_2$ S45W/K46L/Q50W | DKDNVAPRWLISPWGY-$NH_2$ |
| AM(37-52)$NH_2$ S45W/K46W/Q50W | DKDNVAPRWWISPWGY-$NH_2$ |
| AM(37-52)$NH_2$ K46L/S48G/Q50W | DKDNVAPRSLIGPWGY-$NH_2$ |
| AM(37-52)$NH_2$ S45R/K46L/S48G/Q50W | DKDNVAPRRLIGPWGY-$NH_2$ |
| AM(37-52)$NH_2$ S45W/K46L/Q50W/Y52F | DKDNVAPRWLISPWGF-$NH_2$ |
| AM(37-52)$NH_2$ S45W/K46G/Q50W/Y52F | DKDNVAPRWGISPWGF-$NH_2$ |
| AM2/IMD(32-47)$NH_2$ H45W | GRQDSAPVDPSSPWSY-$NH_2$ |
| AM2/IMD(32-47)$NH_2$ S43G/H45W | GRQDSAPVDPSGPWSY-$NH_2$ |
| αCGRP(1-37)$NH_2$ | ACDTATCVTHRLAGLLSRSGGVVKNNFVPTNVGSKAF-$NH_2$ |
| αCGRP(1-37)$NH_2$ N31D/S34P/K35W/A36S **"ss-CGRP"** | ACDTATCVTHRLAGLLSRSGGVVKNNFVPTDVGPWSF-$NH_2$ |
| αCGRP(8-37)$NH_2$ | VTHRLAGLLSRSGGVVKNNFVPTNVGSKAF-$NH_2$ |
| αCGRP(8-37)$NH_2$ N31D/S34P/K35W/A36S | VTHRLAGLLSRSGGVVKNNFVPTDVGPWSF-$NH_2$ |
| αCGRP(27-37)$NH_2$ N31D/S34P/K35W/A36S | FVPTDVGPWSF-$NH_2$ |
| αCGRP(27-37)$NH_2$ N31D/S34P/K35W/A36S/F37P | FVPTDVGPWSP-$NH_2$ |
| αCGRP(27-37)$NH_2$ N31D/S34P/K35W/A36S/F37A | FVPTDVGPWSA-$NH_2$ |
| FITC-AM(37-52)$NH_2$ S45W/Q50W | FITC-Ahx(aminohexanoic acid)-DKDNVAPRWKISPWGY-$NH_2$ |

* All cysteines are in the oxidized disulfide bonded form.

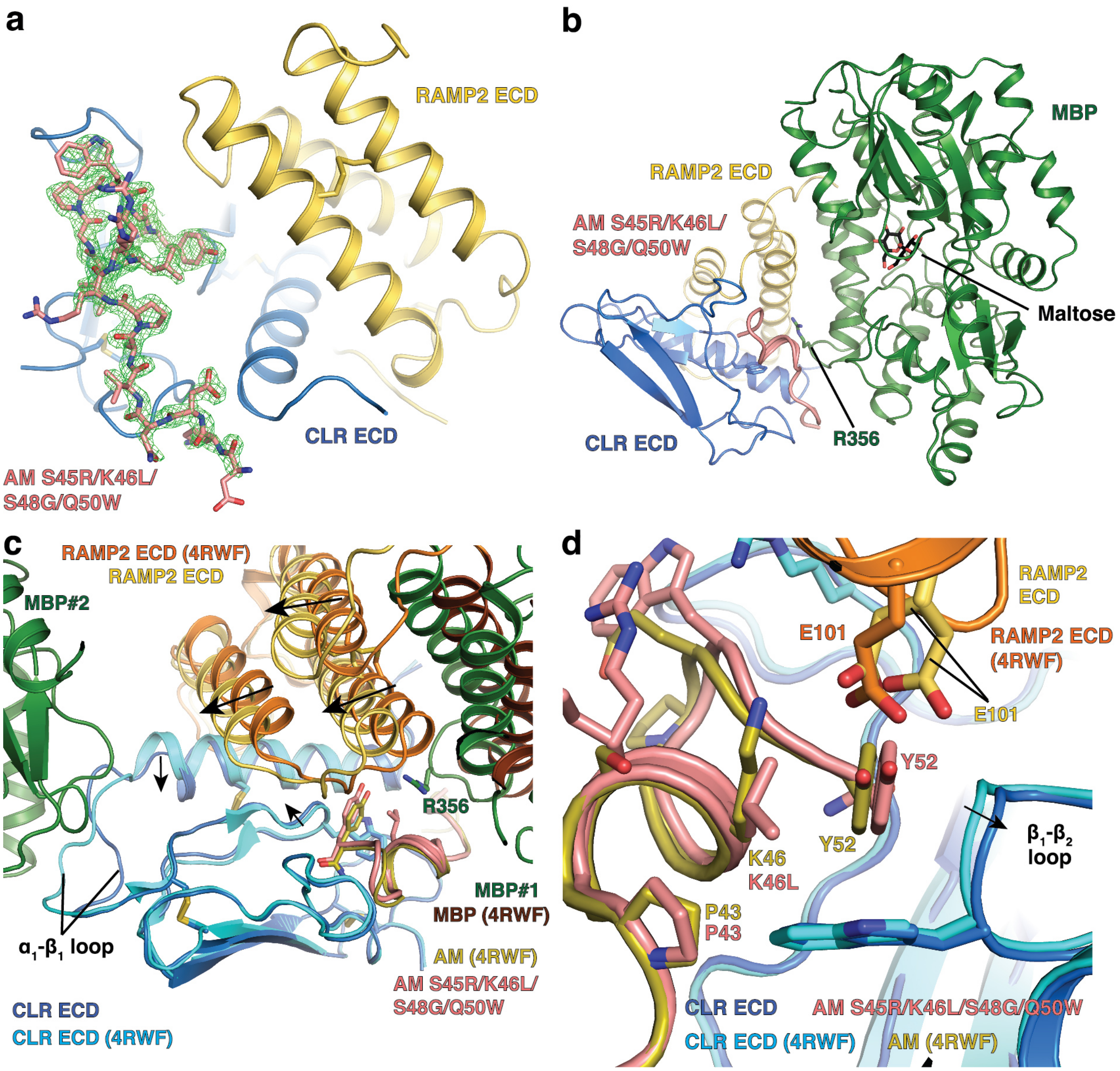

**Supplementary Figure 1. Electron density and crystal packing for the structure of AM S45R/K46L/S48G/Q50W-bound MBP-RAMP2-CLR ECD complex.** (a) $m\mathrm{F_o}$-$D\mathrm{F_c}$ electron density map of peptide contoured at 3σ (green mesh) obtained after rebuilding and refinement of the molecular replacement solution but before peptide modeling. Final AM peptide model shown as reference. A 3Å carve around peptide is shown and MBP is omitted for clarity. (b) Overview of structure with maltose-binding protein (MBP) interactions. R356 of MBP and maltose are shown in stick representation. (c) Superposition of AM-bound MBP-RAMP2-CLR ECD [PDB 4RWF] and AM variant-bound MBP-RAMP2-CLR ECD structures. Structures are aligned based CLR Cα position. MBP#2 is a second MBP molecule from a symmetry mate in AM variant-bound MBP-RAMP2-CLR ECD structure. Arrows indicate changes from AM-bound MBP-RAMP2-CLR ECD structure [PDB 4RWF] to new AM variant structure. (d) Detailed view of peptide-binding pocket shows altered position of $\beta_1$-$\beta_2$ loop. Key residues are shown in stick representation. Two conformations of RAMP2 E101 were modeled. Arrow indicates difference between structures as shown in panel c.

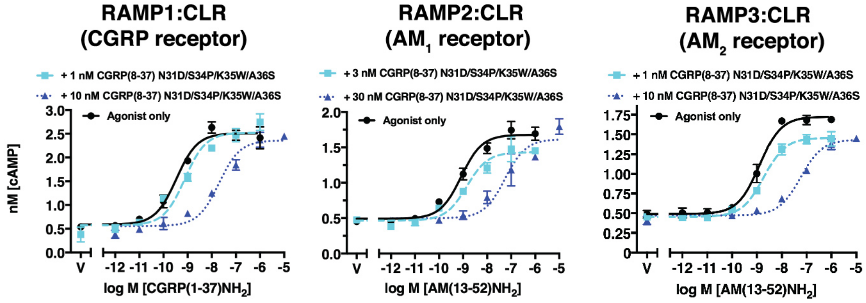

**Supplementary Figure 2. COS-7 cell-based cAMP signaling antagonism assay with simultaneous addition of agonist and antagonist.** Representative concentration-response curves of indicated concentrations of CGRP(8-37)$NH_2$ N31D/S34P/K35W/A36S antagonist using transiently expressed RAMP:CLR receptors in COS-7 cells. Following a 30 minute pre-incubation in the absence of antagonist or agonist, both antagonist and agonist were added simultaneously with continuous exposure to receptors for 30 minutes. Error bars are shown as SD. Data shown are representative from two independent experiments with duplicate technical replicates.

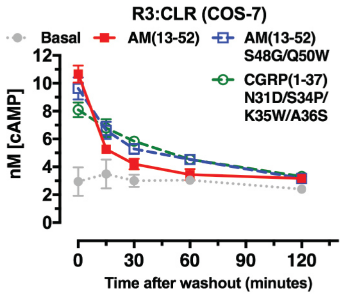

**Supplementary Figure 3. Ligand washout assay for the $AM_2$ receptor (RAMP3:CLR) in COS-7 cells.** Representative cAMP signaling response over time following ligand washout. 100 nM of each peptide agonist was exposed to receptors for 10 minutes before extensive washing with assay buffer (see Methods). Error bars are SD. Data shown are representative from two independent experiments with duplicate technical replicates.

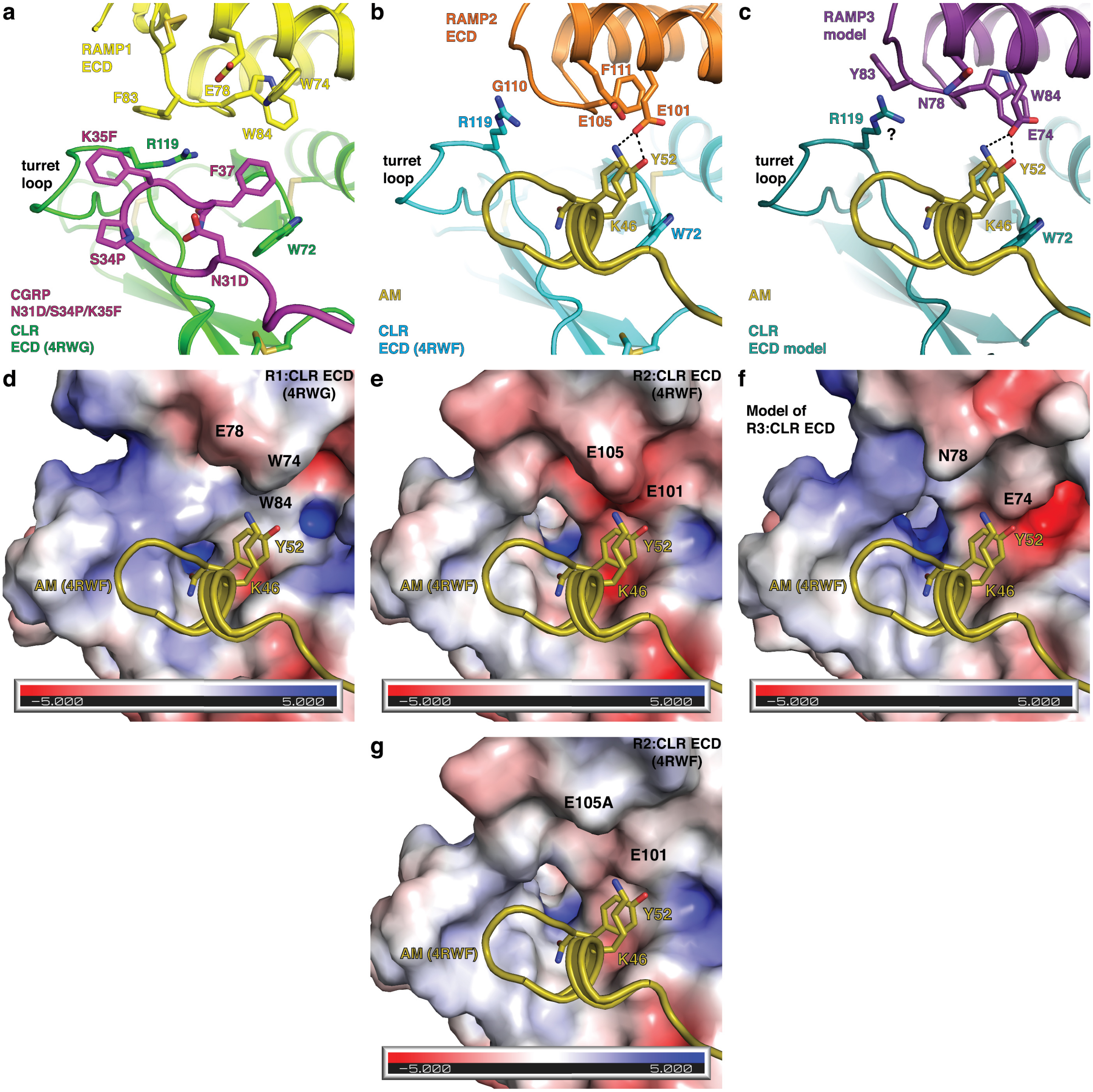

**Supplementary Figure 4. Determinants of AM and CGRP peptide selectivity at RAMP:CLR ECD complexes.** (a-c) Cartoon representation of (a) CGRP N31D/S34P/K35F-bound MBP-RAMP1-CLR ECD [PDB 4RWG], (b) AM-bound MBP-RAMP2-CLR ECD [PDB 4RWG], or (c) modeled AM-bound RAMP3-CLR ECD complex (Booe et al. (2015). *Mol Cell*.). Key residues are shown in stick representation. MBP is omitted for clarity. All colors used are consistent with Figure 1. (d-g) Electrostatic surface representation of AM-bound (d) MBP-RAMP1-CLR ECD complex, (e) MBP-RAMP2-CLR ECD complex, (f) modeled RAMP3-CLR ECD complex, or (g) MBP-RAMP2-CLR ECD complex with modeled E105A mutation. E105A was previously shown to have no effect on AM signaling potency (Booe et al. (2015). *Mol Cell*.). Electrostatic surface was generated in PyMOL with APBS plugin. Surface shown is solvent excluded surface (Connolly surface). All panels were created from the same view in PyMOL.
